# Efficient Homology-directed Repair with Circular ssDNA Donors

**DOI:** 10.1101/864199

**Authors:** Sukanya Iyer, Aamir Mir, Joel Vega-Badillo, Benjamin P. Roscoe, Raed Ibraheim, Lihua Julie Zhu, Jooyoung Lee, Pengpeng Liu, Kevin Luk, Esther Mintzer, Josias Soares de Brito, Philip D. Zamore, Erik J. Sontheimer, Scot A. Wolfe

**Author notes:** These authors contributed equally (Sukanya Iyer, Aamir Mir). Correspondence should be addressed to E.J.S. or S.A.W. Inscripta, Inc., 7060 Koll Center Parkway, Suite 312, Pleasanton, CA 94566, USA.

## Abstract

While genome editing has been revolutionized by the advent of CRISPR-based nucleases, difficulties in achieving efficient, nuclease-mediated, homology-directed repair (HDR) still limit many applications. Commonly used DNA donors such as plasmids suffer from low HDR efficiencies in many cell types, as well as integration at unintended sites. In contrast, single-stranded DNA (ssDNA) donors can produce efficient HDR with minimal off-target integration. Here, we describe the use of ssDNA phage to efficiently and inexpensively produce long circular ssDNA (cssDNA) donors. These cssDNA donors serve as efficient HDR templates when used with Cas9 or Cas12a, with integration frequencies superior to linear ssDNA (lssDNA) donors. To evaluate the relative efficiencies of imprecise and precise repair for a suite of different Cas9 or Cas12a nucleases, we have developed a modified Traffic Light Reporter (TLR) system [TLR-Multi-Cas Variant 1 (MCV1)] that permits side-by-side comparisons of different nuclease systems. We used this system to assess editing and HDR efficiencies of different nuclease platforms with distinct DNA donor types. We then extended the analysis of DNA donor types to evaluate efficiencies of fluorescent tag knock-ins at endogenous sites in HEK293T and K562 cells. Our results show that cssDNA templates produce efficient and robust insertion of reporter tags. Targeting efficiency is high, allowing production of biallelic integrants using cssDNA donors. cssDNA donors also outcompete lssDNA donors in template-driven repair at the target site. These data demonstrate that circular donors provide an efficient, cost-effective method to achieve knock-ins in mammalian cell lines.

## Introduction

RNA-guided Cas9^1–3^ and Cas12a proteins^4, 5^ have provided a facile tool for introducing targeted breaks within genomes. These double-strand breaks (DSBs) can be harnessed to engineer the genome through endogenous DNA repair pathways. Typically, DSBs are precisely repaired via the canonical non-homologous end joining (c-NHEJ) pathway, restoring the original DNA sequence.^4^ However, in the context of a programmable nuclease where DSB generation can reoccur, imprecise DNA repair may produce small insertions and deletions (indels) via c-NHEJ as well as alt-NHEJ pathways.^6^ In contrast to the imprecise nature of these indels, the homology-directed repair (HDR) pathway results in precise rewriting of the genome in a template-dependent manner.^7–9^ HDR is often utilized in the context of programmable nucleases to introduce specific changes to the genome, such as adding fluorescent tags to proteins^10^ or making a precise therapeutic correction to the desired locus.^11–13^ Given the broad utility of this technology for enabling precise insertions into mammalian genomes, several viral and non-viral approaches for the delivery of donor DNA into mammalian cells have been described.^14–17^ The nature of the template employed for HDR is dictated in part by the length of the desired genomic modification. For short insertions (<200 nt), ssDNA oligonucleotides harboring the mutation, as well as flanking homology arms that range from 35-60 nucleotides, are introduced into cells along with Cas9 protein and guide RNA.^15, 18, 19^ When modifications longer than 200bp are desired, double-stranded DNA (dsDNA) templates such as plasmids or PCR products are typically used as donor templates. However, these double-stranded templates are often associated with high cellular toxicity and off-target integration events.^20^ As an alternative to using dsDNA templates as donors for HDR, long ssDNA templates have been reported to have low cytotoxicity and high efficiencies of targeted integration at the site of interest.^21, 22^ Consequently, there is considerable interest in developing methods to generate long ssDNA templates to serve as donors for making targeted insertions in mammalian cells. Several recent examples include asymmetric PCR, “Strandase” enzyme-mediated removal of one strand of a linear dsDNA template [Takara Bio USA (catalogue number 632644)], use of pairs of nicking endonucleases followed by gel extraction of resulting ssDNA [Biodynamics Laboratory Inc. (catalogue number DS615) and reverse transcription (RT)-based approaches to generate ssDNA.^21–24^ Most of these approaches require expensive and time-consuming purification steps to ensure complete removal of truncated ssDNA products. With RT-based approaches in particular, it is challenging to generate accurate ssDNA donors longer than 3-4 kb, especially in large molar quantities, because of the lack of proofreading activity and the limited processivity of reverse transcriptase enzymes.

As an alternative to these *in vitro* approaches, we explored the use of circular ssDNA (cssDNA) produced from phagemids as templates for HDR-mediated integration of DNA cassettes. Phagemid vectors have been used to generate ssDNA templates for site-directed mutagenesis^25^, DNA nanotechnology and DNA origami^26^, phage display technology for protein engineering^27^ and as templates for transcription in cell-free systems.^28^ However, to our knowledge, their use as donors for achieving targeted integration of DNA in mammalian cells has not been evaluated.

Here, we show that phagemid-derived cssDNA can be used to insert sequences efficiently and precisely in mammalian cells. We further compared HDR efficiencies obtained with phagemid-sourced cssDNA to those of linear ssDNAs (lssDNAs) generated using a RT-based method^22^ and a streptavidin affinity purification approach with asymmetrically biotinylated PCR amplicons.^29^ To this end, we utilized a redesigned traffic light reporter system to evaluate HDR efficiencies for different forms of donor templates (plasmids, lssDNAs and cssDNAs) when used in conjunction with SpyCas9 or three different Cas12a effectors delivered as ribonucleoproteins (RNPs) in HEK293T and K562 cells. We then compared knock-in yields of linear and circular ssDNA donor templates containing fluorescent reporter tags at four different endogenous sites in the human genome. Finally, we demonstrated the ability of circular ssDNA templates to create biallelic integration of a reporter cassette in different cell lines. Overall, our data show broad utility of cssDNA as donors for genome engineering applications.

## Results

### Generating linear and circular ssDNA templates for HDR in mammalian cells

To address the challenges associated with long ssDNA donor production, we investigated a number of different approaches for generating ssDNA donors, as well as the relative efficiencies of HDR when using the resulting ssDNA products. While most efforts to generate ssDNA donors have focused on linear molecules, we explored the properties of circular ssDNAs as donors for HDR. Phagemids are chimeric vectors that contain plasmid and bacteriophage origins of replication. Upon superinfection of the host bacteria with helper phage to supply the phage DNA replication machinery, one strand of the phagemid vector is packaged into bacteriophage particles and extruded into the media from whence circular ssDNA can be purified^30^ (Supplementary Fig. S1A). Although a standard protocol to purify ssDNA from phagemids yielded reasonable quantities of DNA, we observed the presence of contaminating *Escherichia coli* genomic DNA in the ssDNA preparation, as reported previously.^31^ To remove contaminating *E. coli* genomic DNA in preparation for donor DNA transfection into mammalian cells, we modified a purification protocol described by Viera and Messing^30^, where we incorporated a DNase I digestion step prior to bacteriophage uncoating and subsequently purified the cssDNA using an anion exchange column.

To provide a benchmark for aspects of donor DNA production and direct comparison of HDR rates in mammalian cells, we also evaluated two methods for generating lssDNA templates. First, lssDNA was generated using a published RT method (T-lssDNA) in which cDNA is generated by a processive reverse transcriptase such as TGIRT-III.^32^ RT-based approaches (Supplementary Fig. S1B) can be effective for generating ssDNA donors up to 3.5 kb in length.^21, 22, 33, 34^ However, the reverse transcriptase enzymes used for generating linear ssDNA generally lack proofreading activity^35^, which makes the fidelity of the resulting template a concern.^24^ In addition, these enzymes often generate truncated ssDNA products (Supplementary Table S1) and yields of full-length ssDNA products, particularly for templates with stable secondary structures, have been found to be compromised.^36^ As an alternative to RT-based methods, we reasoned that ssDNA templates generated from asymmetrically biotinylated PCR products would produce longer ssDNA templates with higher sequence fidelity. Accordingly, we utilized an approach to generate ssDNA templates using biotin-based affinity purification of ssDNA (B-lssDNA) by exploiting the biotin-streptavidin interaction. In this method, one PCR primer used for donor amplification is biotinylated, which allows the resulting PCR product to be strand-specifically bound to streptavidin-coated beads. Subsequently, the DNA strands are separated by alkaline denaturation and the non-biotinylated strand is isolated and used as a donor for HDR (Supplementary Fig. S1C). SsDNA templates generated by all these methods were treated with S1 nuclease to confirm the single-stranded nature of the templates generated (Supplementary Fig. S1D). Overall, while all three approaches yielded ssDNA up to at least 3,300 bases in length, the phagemid-based approach proved to be most economical while also generating large quantities of full-length ssDNA for use as HDR templates (Table 1).

**Table 1:**
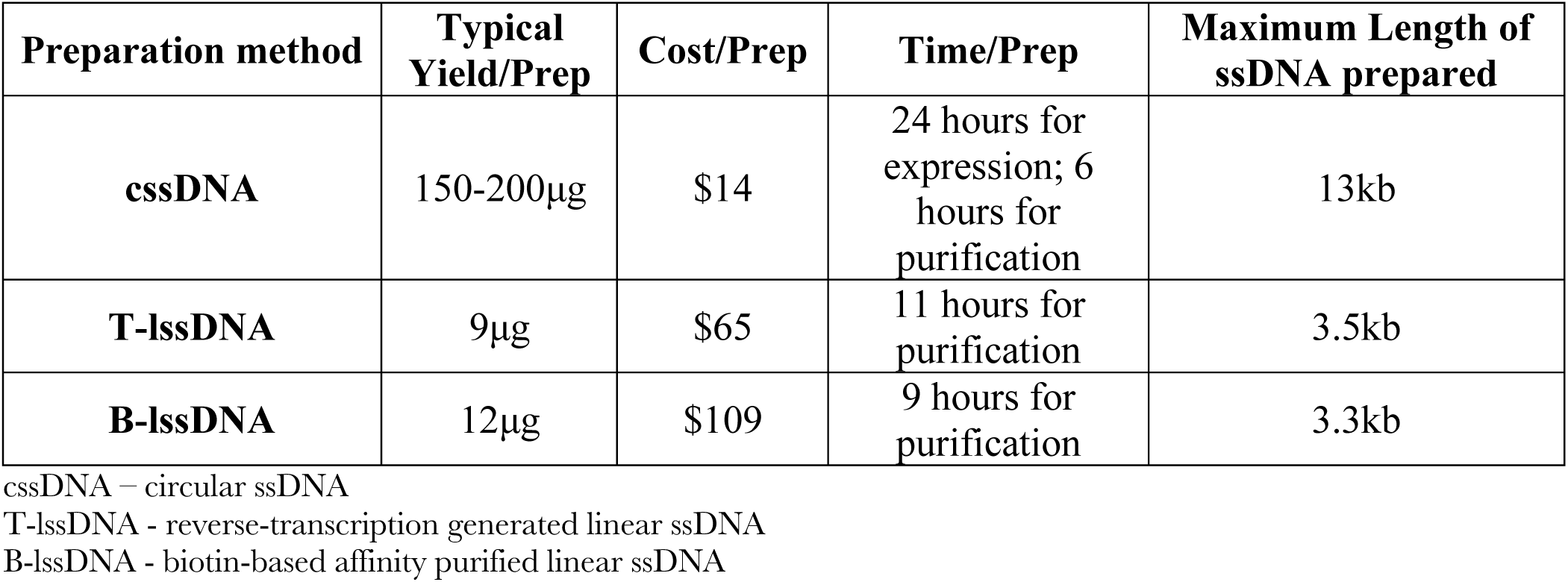
Features of different ssDNA preparation methods.

### Traffic Light Reporter Multi-Cas Variant 1 (TLR-MCV1): a system to evaluate genome-editing efficiency by multiple nucleases

Previously, Certo *et al.* described a traffic light reporter (TLR) system that provides positive fluorescence readouts for both error-prone DSB repair as well as precise HDR repair.^37^ It consists of a tandem expression cassette consisting of a “broken” GFP coding sequence followed by an out-of-frame mCherry cassette (Figure 1A). The GFP sequence is disrupted by an insertion harboring various nuclease target sites to initiate DSB formation. DSB repair by pathways such as NHEJ can result in insertions or deletions (indels) that place the downstream mCherry coding sequence in frame for productive translation (+1 frameshift). In addition, precise HDR repair of the locus can be evaluated by co-delivering a truncated GFP donor repair template with a nuclease, which will restore GFP expression while leaving the mCherry coding sequence out of frame. The fraction of GFP- and mCherry-positive cells can be rapidly measured using flow cytometry to determine editing outcomes as a function of the nuclease and donor DNA composition. We redesigned the original TLR reporter to incorporate target sites for several currently characterized nucleases (Figure 1A) by introducing protospacer adjacent motifs (PAMs) belonging to Cas9/Cas12a orthologs from *Streptococcus pyogenes* (SpyCas9)^38, 39^, *Neisseria meningiditis* (Nme1Cas9 and Nme2Cas9)^40–42^, *Campylobacter jejuni* (CjeCas9)^43–45^, *Staphylococcus aureus* (SauCas9)^46^, *Geobacillus stearothermophilus* (GeoCas9)^47^, *Lachnospiraceae* bacterium ND2006 (LbaCas12a)^48^, *Acidaminococcus* sp. (AspCas12a)^48^ and *Francisella novicida* (FnoCas12).^49^ For several of the Cas9 orthologs (SpyCas9, Nme1Cas9, CjeCas9 and SauCas9), DSB formation can be targeted to the exact same position. We also incorporated a second SpyCas9 target site on the opposite strand such that both SpyCas9 target sites will produce a DSB at the same position. Similarly, the Cas12a orthologs have overlapping PAMs in the incorporated target site and therefore will generate staggered cuts within the same region. All of these target sites were combined into a sequence framework that lacks stop codons in the +1 reading frame to enable mCherry expression following the induction of a suitable indel. Hence, our updated reporter (TLR-MCV1) provides a useful platform for direct comparison of genome editing properties of the major RNA-guided genome editing tools described to date.

**Fig. 1.**
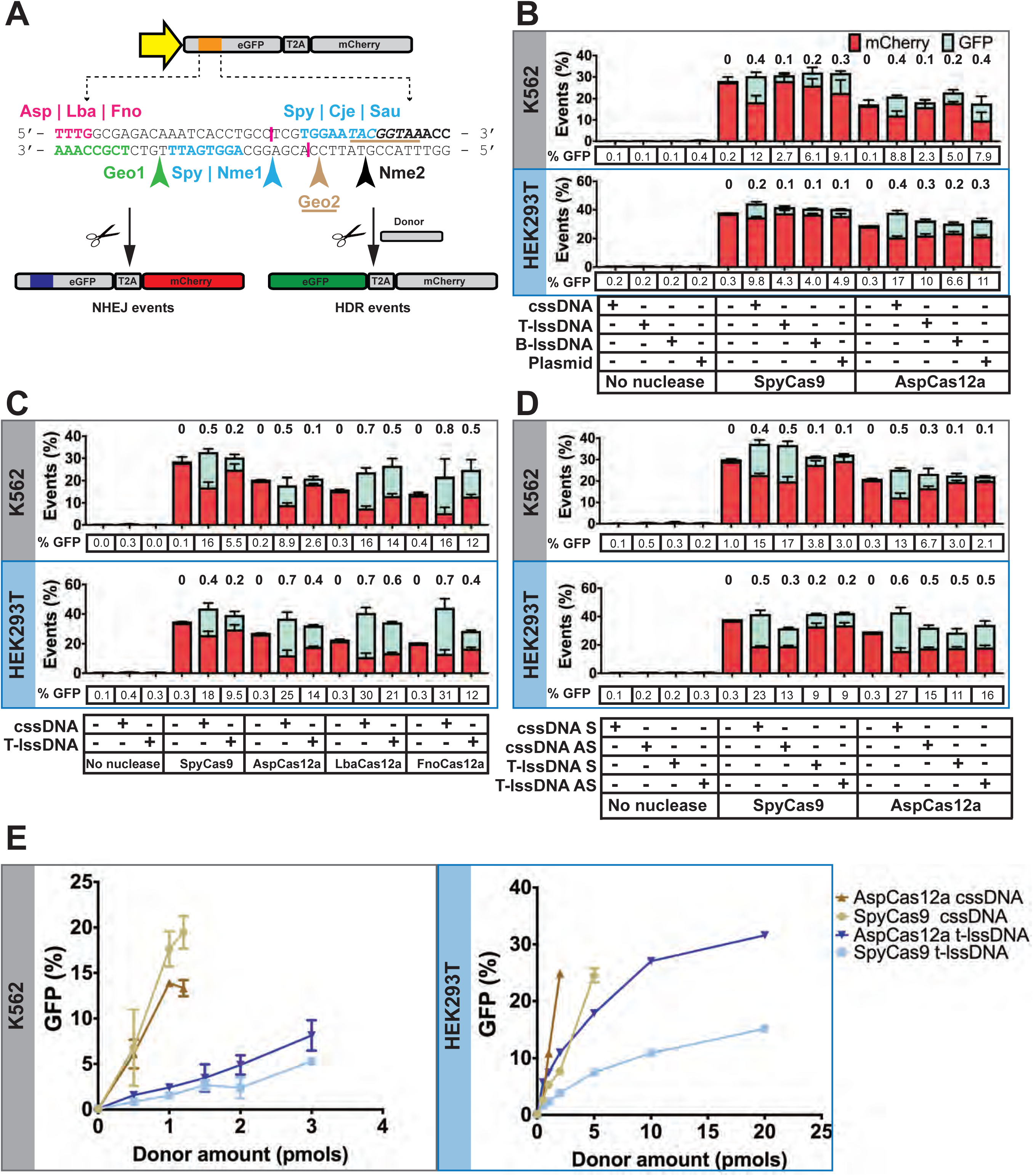
Comparisons of the activity of different DNA donors in homology-directed repair using the Traffic Light Reporter Multi-Cas Variant 1 (TLR-MCV1) cassette in human cells. **(A)** The schematic depicts the TLR-MCV1 system showing the SFFV promoter driving the expression of GFP and mCherry, separated by a ribosome-skipping T2A signal. The yellow arrow depicts the SFFV promoter driving the expression of the GFP-T2A-mCherry cassette. The orange line indicates the insertion containing target sequences for different Cas effectors, the sequence of which is shown below the schematic of TLR-MCV1. Sequences and arrows in blue indicate overlapping PAMs and a common cut site associated with SpyCas9, Nme1Cas9, CjeCas9 and SauCas9. The bolded black sequence and black arrow depict the Nme2Cas9 PAM and cut site respectively. Magenta text shows PAMs associated with Cas12a effectors, and their approximate cut sites are shown by magenta lines. The PAMs associated with Geo1Cas9 and Geo2Cas9 are highlighted in green and tan text, respectively. The cut sites for these two Cas9s are shown by green and tan arrows, respectively. DSBs at any of the sites of these may be imprecisely repaired via the NHEJ pathway resulting in mCherry expression (shown on the left) if repair results in (+1 frameshift) productive translation. In the presence of donor, HDR-mediated correction of “broken” GFP region results in restoration of GFP expression (shown on the right). **(B)** Efficacy of distinct DNA templates in driving HDR. The graph depicts the percentage of mCherry- and GFP-positive cells obtained after co-delivery of SpyCas9 or AspCas12a RNP with cssDNA, T-lssDNA, B-lssDNA or plasmid DNA repair templates into TLR-MCV1 K562 cells (upper grey box) and TLR-MCV1 HEK293T cells (lower blue box). Numbers above the bars indicate ratios of GFP-positive (shown in cyan) to total indel events [mCherry-positive (shown in red) + GFP-positive cells]. Bars represent the mean from three independent biological replicates and error bars represent the standard error of the mean (s.e.m.). **(C)** Comparison of cssDNA- and T-lssDNA-mediated HDR efficiency upon treatment of TLR-MCV1 cells with distinct Cas effectors. The graph depicts the percentage of mCherry- and GFP-positive cells obtained after co-delivery of SpyCas9, AspCas12a, LbaCas12a or FnoCas12a with cssDNA and T-lssDNA DNA repair templates into TLR-MCV1 K562 cells (upper grey box) and TLR-MCV1 HEK293T cells (lower blue box). Numbers above the bars indicate ratios of GFP-positive (shown in cyan) to total indel events (mCherry-positive + GFP-positive). Bars represent the mean from three independent biological replicates and error bars represent the standard error of mean (s.e.m.). **(D)** Effect of cssDNA and T-lssDNA donor orientation on HDR efficiency. The graph depicts the percentage of mCherry- and GFP-positive cells obtained after co-delivery of SpyCas9-1 or AspCas12a (targeting the same strand) with sense (S) and antisense (AS) strand cssDNA and T-lssDNA DNA repair templates into TLR-MCV1 K562 cells (upper grey box) and TLR-MCV1 HEK293T cells (lower blue box). Numbers above the bars indicate ratios of GFP-positive (shown in cyan) to total indel events (mCherry-positive + GFP-positive). Bars represent the mean from three independent biological replicates and error bars represent s.e.m. **(E)** Dose dependence of cssDNA and T-lssDNA donor template-mediated HDR efficiency. The graph shows the percentage of GFP-positive cells as a function of increasing cssDNA and T-lssDNA donor DNA in the presence of SpyCas9 and AspCas12a proteins in TLR-MCV1 K562 cells (left) and HEK293T cells (right). Bars represent the mean from three independent biological replicates and error bars represent s.e.m.

A single copy of TLR-MCV1 was introduced into HEK293T and K562 cells by lentiviral transduction. Using plasmid transfections of HEK293T cells to introduce the nucleases, guide RNA (listed in Supplementary Table S2) and a plasmid donor template (pCVL-SFFV-d14GFP-Donor; Supplementary Table S3), we observed that all the Cas9/Cas12a sites can be targeted by the cognate nucleases to induce precise and imprecise genome editing in mammalian cells (Supplementary Fig. S2A). The two GeoCas9-expressing plasmids produced inefficient editing, which may be due to suboptimal codon usage, or to GeoCas9’s preference for higher temperatures, or both.^47^ We also performed a dose-dependence analysis to test the potency of different nucleases (Supplementary Fig. S2B). SpyCas9 was found to be the most potent nuclease for the production of frameshifts that restore mCherry expression.

### Circular ssDNA donors outperform linear ssDNA donors for HDR

TLR-MCV1 provides an ideal system for direct comparisons of different DNA donor architectures since both the NHEJ and HDR efficiencies can be measured using different Cas nucleases at the same locus. To create DSBs in cells, delivery of Cas9 or Cas12a RNPs has gained favor because these complexes can be readily electroporated into a wide variety of cell types.^50–53^ Furthermore, due to their rapid turnover in cells, Cas9/Cas12a RNPs display lower off-target activity than other delivery modalities without compromising on-target editing activity, thereby significantly improving the specificity of targeted genomic modifications.^51, 54^ Delivery of SpyCas9 protein complexed with its guide RNA (SpyCas9 RNPs), or each of the three Cas12a orthologs as RNPs, proved highly effective at editing the TLR-MCV1 reporter, with indel efficiencies greater than 70% achieved as measured by TIDE^55^ (Supplementary Fig. S3). Next, we tested different types of ssDNA donors or a plasmid donor with SpyCas9 and AspCas12a RNPs. As shown in Figure 1B, cssDNA elicited higher HDR efficiencies relative to equimolar quantities of linear ssDNA donors or the plasmid donor in both K562 and HEK293T cells. Using cssDNA, we achieved a statistically significant ~2-fold increase in HDR yields compared to lssDNA (Supplementary Table S4). This was true for both SpyCas9 and AspCas12a-based editing. CssDNA also achieved higher GFP integration efficiencies in comparison to plasmid donors in both K562 and HEK293T cells. Notably, we did not observe a significant difference between T-lssDNA and B-lssDNA donor efficiency in K562 cells (*p* = 0.0797), indicating that lssDNAs generated using two different approaches were largely indistinguishable once generated and purified (Supplementary Table S4). There was a statistically significant difference (*p* = 0.03) between T-lssDNA and B-lssDNA when tested in HEK293T cells with AspCas12a. However, the increase shown by T-lssDNA relative to B-lssDNA is modest (<4%). Overall, among the different forms of DNA templates tested, cssDNA realized the highest HDR efficiencies.

The improved efficiency of knock-in using cssDNA may be due to increased exonuclease protection afforded by the circular nature of the ssDNA. To test this hypothesis, we circularized the lssDNA by splint-mediated ligation and tested this circularized form in TLR-MCV1 cells (Supplementary Fig. S4A). Circularization of linear ssDNA resulted in significant (*p* < 0.0001) enhancement of HDR relative to the unligated precursor in both the cell lines (Supplementary Fig. S4B, Supplementary Table S4) and comparable efficiencies to those observed with phagemid-derived cssDNA donors. This is consistent with previous studies that demonstrated improved function of end-protected nucleic acids in various cell types.^56^

### Cas12a nucleases produce superior HDR yields at the TLR-MCV1 locus

Cas12a-based genome editing has been reported to achieve increased HDR, relative to SpyCas9, since it generates 5’ overhangs and more rapidly releases the PAM-distal DNA end following cleavage.^57^ As shown in Figure 1B, in HEK293T cells the HDR efficiency as a fraction of total editing ([GFP/(GFP + mCherry)], referred to hereafter as the “HDR ratio”) with all the donors tested was higher for AspCas12a compared to SpyCas9. By contrast, we did not observe increases in the HDR ratio of editing with AspCas12a compared to SpyCas9 in K562 cells. To explore this observation further, we tested different orthologs of Cas12a with lssDNA and cssDNA donors. Since we had previously observed no substantial difference between B-lssDNA and T-lssDNA in HDR efficiency at the TLR-MCV1 locus, we only included T-lssDNA for the subsequent comparisons in TLR-MCV1-related experiments. Efficacy of different SpyCas9 and Cas12a nucleases for driving HDR showed cell-line-specific differences. The LbaCas12a and FnoCas12a variants yielded higher HDR ratios relative to SpyCas9 (Figure 1C) in both HEK293T and K562 cells. With AspCas12a, however, while HDR ratios are increased in HEK293T cells, a similar increase in HDR ratios was not observed in K562 cells. In HEK293T cells, SpyCas9 supported HDR percentages of 18% and 9.5% with cssDNA and lssDNA donors, respectively (Figure 1C, lower panel). Cas12a orthologs increased HDR percentages to 25-31% with cssDNA template and 12-21% with linear ssDNA donor. Among the Cas12a orthologs tested, LbaCas12a and FnoCas12a showed higher HDR ratios compared to AspCas12a with cssDNA. In K562 cells, the same trends were generally observed, with the exception of editing efficiencies for AspCas12. In K562 cells, the HDR ratio increased from 0.5 with SpyCas9 to 0.7-0.8 with LbaCas12a and FnoCas12a when using the cssDNA donor (Supplementary Fig. S5). Thus, in these cells the HDR pathway was predominantly being harnessed for DSB repair during Cas12a-mediated genome editing with the cssDNA donor. The overall HDR ratio with the linear ssDNA donor increased to approximately 0.5 with LbaCas12a and FnoCas12a (Supplementary Fig. S5). However, AspCas12a did not show similar enhancements in HDR ratio in K562 cells. Taken together these results indicate that Cas12a orthologs may be superior for template-dependent HDR genome editing when compared to SpyCas9.

### The effect of donor orientation is dependent on cell type and nuclease identity

There are conflicting reports in the literature regarding the effect of DNA strand orientation on HDR efficiencies. A bias in HDR efficiency towards ssDNA donors that have the same sequence as the target strand (i.e. the strand base-paired to the SpyCas9 RNA guide) has been reported.^18, 58^ However, others have not observed a significant effect of strand orientation on HDR efficiency.^15, 57, 59^ To examine strand-specific donor bias in HDR efficiencies in TLR-MCV1 cells, we generated target-strand-complementary (sense) and non-target-strand-complementary (antisense) ssDNA donors for both linear and circular DNAs and electroporated them along with SpyCas9 and AspCas12a RNPs. For both effectors, the guide RNA was complementary to the antisense strand of the TLR-MCV1 reporter. In K562-TLR-MCV1 cells, there were no significant differences between sense and antisense ssDNA donors except in the case of AspCas12a and cssDNA donors (Figure 1D). For this effector/donor combination, there was about a 2-fold increase in HDR efficiency with the sense cssDNA donor relative to antisense cssDNA donor. On the other hand, electroporated HEK293T cells exhibited higher HDR yields (*p* < 0.008) with sense cssDNA donors when used with both SpyCas9 and AspCas12a. The increase in the HDR efficiency with sense cssDNA relative to antisense cssDNA was 7% and 13% when cells were electroporated with SpyCas9 and AspCas12a, respectively. To examine if the two different guide orientations relative to the coding region of the TLR-MCV1 sequence influence the ssDNA donor orientation preference for HDR for SpyCas9 in K562 cells, we electroporated cssDNA and lssDNA donors that were complementary to the TLR-MCV1 sense or the antisense strand, in combination with guide RNAs that were likewise complementary to either TLR-MCV1 target site strand (Supplementary Fig. S6A). We did not observe any significant differences in HDR efficiency as a function of relative guide/donor strand orientation (Supplementary Fig. S6B). Overall, while there are nuclease- and cell-type-specific differences HDR efficiencies, the relative orientation of the donor does not have a consistent impact on HDR-based editing. This is consistent with previously described ssDNA donor strand biases in HDR efficiencies, which are generally locus- and cell type-specific^19^.

### Circular ssDNA donors are more potent than lssDNA donors for HDR

We reasoned that the higher nuclease stability of cssDNA donors may improve the potency of cssDNA compared to lssDNA donors. To test this hypothesis, cells were electroporated with increasing amounts of ssDNA donors while keeping the amount of SpyCas9 or AspCas12a RNPs constant (Figure 1E). In K562 cells, the HDR yields peaked around 1pmol of cssDNA for both SpyCas9 and AspCas12a. We also observe severe apparent DNA toxicity at higher donor DNA concentrations (>1 pmoles of cssDNA) resulting in reduction of HDR efficiencies. Since overall cssDNA templates are about 4-5 times longer due to the presence of the phagemid sequence elements, it’s likely that DNA toxicity is associated with the total mass of DNA delivered instead of moles of DNA templates electroporated. Even so, the lssDNA donor did not perform as well as the cssDNA donor in stimulating HDR even at the highest concentration that was tested in K562 cells. The highest HDR efficiency observed for the lssDNA was about 5% with SpyCas9 and 7% with AspCas12a which is four and two times lower than what was achieved with the cssDNA donor and SpyCas9 and AspCas12a respectively. These results were also mirrored in HEK293T cells, where the cssDNA donor was more potent compared to lssDNA donor. With AspCas12a, cssDNA reached saturation at around 2 pmols, whereas 5 pmols was needed to achieve the same effect with SpyCas9. Above these donor DNA levels, we observed a drop in HDR efficiencies, presumably due to DNA toxicity. The lssDNA donor performed poorly with SpyCas9 since the percentage of GFP-positive cells with 20 pmoles of donor was still ~10% lower despite using 3-fold more moles of donor. The lssDNA performed better with AspCas12a where HDR efficiencies of ~30% were achieved with 20 pmoles of lssDNA donor. However, to achieve the same HDR yields, 5-fold more moles of lssDNA was needed compared to cssDNA donor. Hence, cssDNA is more potent than lssDNA for HDR and its effect is further enhanced when employing AspCas12a as the nuclease. Collectively, the TLR-MCV1-based experiments reveal that cssDNA donors are more efficient at promoting HDR repair compared to lssDNA donors.

### Circular ssDNA donors provide efficient templates for fluorescent tagging of endogenous proteins

For many functional genomic studies and gene therapy applications, targeted insertion of long DNA cassettes into endogenous loci is desirable. Most studies aimed at making targeted insertions of long DNA cassettes employ plasmid donors to provide the template for precise insertion.^10^ However, plasmid donors can be toxic to target cells, which makes insertion of long DNA cassettes an inefficient process in most cell types.^16^ To test the suitability of cssDNA for integrating larger inserts, we chose four endogenous genes in the mammalian genome based on the work of Roberts *et al*.^10^ and He *et al*.^60^ to make targeted insertions of fluorescent proteins (Figure 2A). SpyCas9 RNPs were complexed with chemically synthesized guide RNAs (listed in Supplementary Table S2) with terminal modifications to enhance intracellular stability. Electroporation of RNPs in the absence of donor DNA into HEK293T cells yielded 80-93% indels at the four sites as measured by TIDE analysis^55^ (Supplementary Fig. S7), indicating efficient SpyCas9 editing of each endogenous locus. It should be noted that while guides targeting *ACTB*, *TOMM20* and *GAPDH* loci are complementary to the sense strand, the *SEC61B* targeting guide is complementary to the antisense strand. To evaluate the relative efficiency of targeted insertion by cssDNA and lssDNA, we tagged three endogenous ORFs (*SEC61B*, *TOMM20* and *ACTB*) via a direct fusion of mEGFP (Figure 2A). At the *GAPDH* locus, we inserted an IRES-EGFP cassette to facilitate separate expression of both gene products from the modified locus.^60^ To evaluate the impact of the donor cassette sequence composition on HDR efficiency, the GFP tag was replaced with a red fluorescence tag (dTomato/iTag RFP) in a corresponding donor set. Phagemid-derived cssDNA or T-lssDNA donors encoding the fluorescence tag flanked by 1kb homology arms were electroporated into K562 and HEK293T cells along with SpyCas9 RNPs, after which GFP- or RFP-positive cells were measured by flow cytometry to estimate the HDR-based recoding efficiency at each site of interest.

**Fig. 2.**
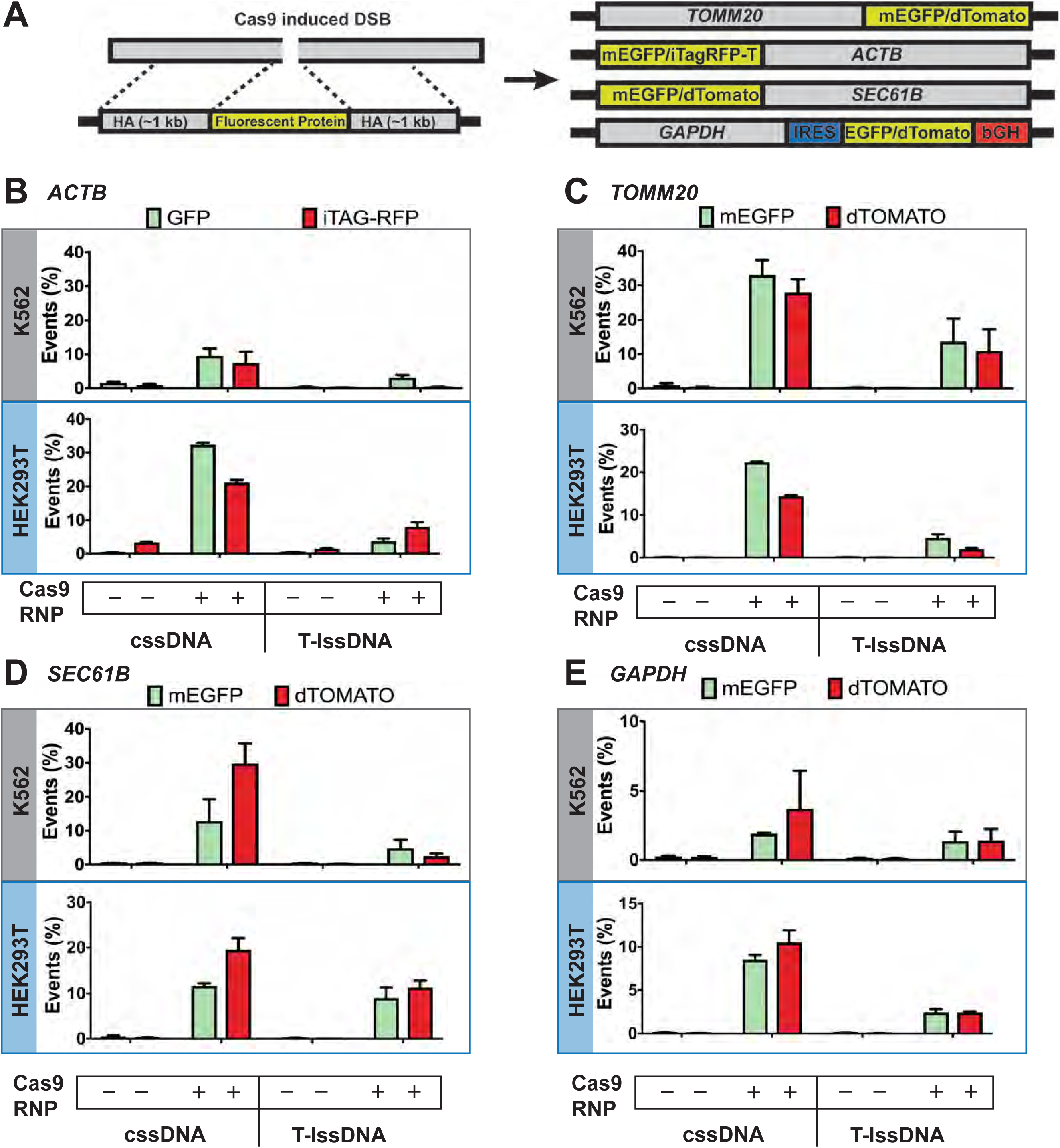
Comparisons of DNA donors in homology-directed repair of endogenous genomic loci in human cells. **(A)** Schematic of fluorescent protein tagging. The left panel shows a schematic of a genomic region containing the SpyCas9 target site and also depicts the design of a donor template containing the fluorescent protein of interest flanked by homology arms (HA). The right panel shows a schematic of each target genomic locus and the arrangement of the fluorescent tag (EGFP, dTomato or iTagRFP-T) following integration. In cases of donors delivered to fluorescently tag the *GAPDH* locus, the fluorescent tag is preceded by an IRES (internal ribosome entry site) and followed by a bovine growth hormone (bGH) polyadenylation sequence. **(B-E)** Bar graphs displaying the percentages of fluorescent cells obtained upon co-delivery of 20 pmoles of SpyCas9 complexed with 25 pmoles of guide RNA targeting the **(B)** *ACTB*, **(C)** *TOMM20*, **(D)** *SEC61B*, or **(E)** *GAPDH* locus with or without cssDNA or T-lssDNA as a donor template. Bars represent the mean from three independent biological replicates and error bars represent s.e.m.

Collectively at all the loci tested, cssDNA resulted in a significantly higher frequency of functional tag integration compared to the linear T-lssDNA (Figure 2B-E; significance values computed in Supplementary Table S4). Interestingly, although GFP and iTagRFP and dTomato fusion tags have coding sequences of similar length, we observed higher integration efficiency with GFP cssDNA donor at the *ACTB* and *TOMM20* locus, especially in HEK293T cells, indicating that donor cassette composition may modestly influence integration efficiency in a cell type- and locus-specific manner. Similarly, at the *SEC61B* locus, cssDNA mediated integration of the dTomato tag was higher than what was achieved with T-ssDNA in both K562 cells and HEK293T cells (Figure 2D). As expected, we did not observe significant differences in donor integration efficiencies between T-lssDNA and B-lssDNA donors, although variability in the efficacy was observed depending on the target site, donor composition and cell type (Supplementary Fig. S8). As with TLR-MCV1, we observed cell-type- and site-specific differences in editing efficiencies with different cssDNA donor orientations, but there was no consistent trend that defined a preferred combination of target site and donor template strand (Supplementary Fig. S9). Collectively, while we observe cell type-, locus- and donor DNA sequence- and orientation-dependent variability in DNA integration efficiencies, our results show the increased potency of cssDNA templates for tagging proteins at various endogenous genomic loci in comparison to lssDNA templates.

### Circular ssDNA can effectively drive biallelic tagging of endogenous proteins

Biallelic tagging of a target gene is often desirable for functional genomics studies, but this outcome is often hampered by low HDR efficiency. Since we observed high yields of integration with cssDNA, we tested the ability of cssDNA to support biallelic integration at various endogenous sites. To distinguish between monoallelic and biallelic integration, we electroporated equimolar amounts of cssDNA donors containing green and red fluorescent tags along with the appropriate SpyCas9 RNP into cells and measured fluorescence in these cells using flow cytometry. The majority of labeled cells expressed a single green or red fluorescent tag (Figure 3). Encouragingly, for *ACTB, TOMM20* and *SEC61B* loci, 17-26% of fluorescent cells were tagged with both green fluorescent and red fluorescent proteins, indicating biallelic integration of reporter tags at these sites (Supplementary Fig. S10). Negligible levels of biallelic integration were observed at the *GAPDH* locus, likely due to lower overall HDR efficiencies at this locus, which could reflect toxicity associated with tagging GAPDH, an essential housekeeping protein.

**Fig. 3.**
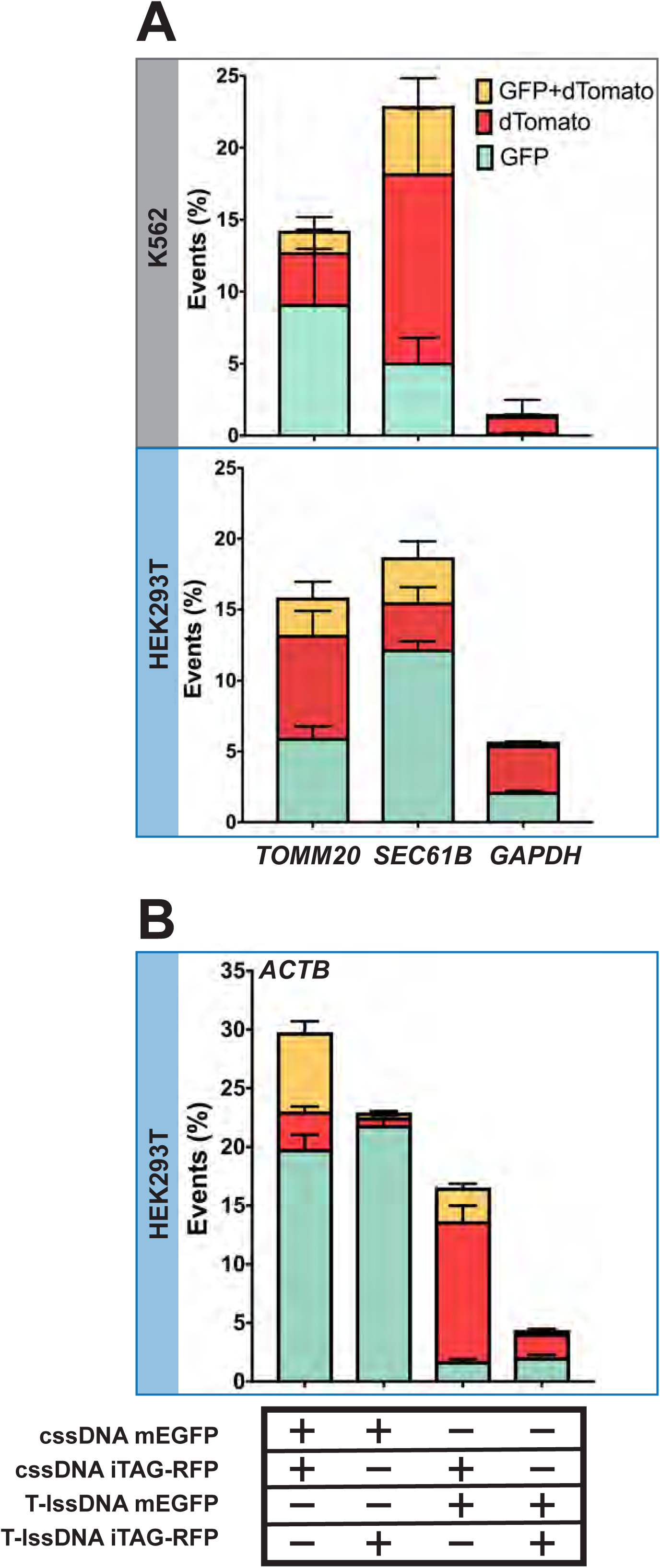
Biallelic tagging of endogenous proteins using two different cssDNA donor templates. **(A)** The graph shows the percentage of fluorescent cells tagged with GFP (shown in cyan), dTomato (shown in red) or both (shown in yellow) at each locus (*TOMM20*, *SEC61B* or *GAPDH*) in K562 cells (top panel) and HEK293T cells (bottom panel). 20 pmol SpyCas9 RNPs were co-delivered with 0.5 pmol of each cssDNA templates. Bars represent the mean from three independent biological replicates and error bars represent s.e.m. **(B)** Competition between cssDNA and lssDNA templates as donors for HDR. The graph shows the percentage of cells tagged with GFP (shown in cyan), iTAG-RFP (shown in red) or both GFP and iTAG-RFP (shown in yellow) at the *ACTB* locus. Bars represent the mean from three independent biological replicates and error bars represent s.e.m.

To further compare the efficiency of fluorescent tag integration at the genetic loci of interest using cssDNA and lssDNA, we set up a competition assay and tested different combinations of cssDNA and lssDNA donors for their abilities to insert reporter tags at the *ACTB* locus (Figure 3B). We observed robust biallelic tagging when cssDNA donors encoding GFP and iTAGRFP tags were cotransfected in both HEK293T cells (Figure 3B) and K562 cells (Supplementary Fig. S11). Interestingly, when cssDNA was combined with an equimolar quantity of lssDNA to perform the knock-ins, we observed 30-fold higher RFP signal over GFP signal when RFP-encoding cssDNA was co-introduced with GFP-encoding lssDNA. Conversely, the combination of GFP-encoding cssDNA with RFP-encoding lssDNA yielded 10-fold more GFP-positive than RFP-positive cells. Overall these results confirm that cssDNA is more efficient than lssDNA as an HDR donor in cultured human cells and is effective for generating biallelic insertions of extended coding sequence into the genome.

## Discussion

For most cellular applications, non-viral methods for the delivery of a donor DNA template are employed to achieve targeted DNA insertion at a locus of interest, owing to the ease of template production. Most previous non-viral approaches have used oligonucleotides (ODNs), plasmids or linear dsDNAs as the donor DNA template.^10, 32, 61–63^ More recently, long lssDNA templates have been demonstrated to provide advantages over dsDNA by both reducing toxicity to cells and increasing HDR efficiency of the DNA donor cassette.^21, 22, 24^ Enzymatic methods adopted for generating long ssDNAs have permitted the knock-in of gene segments such as fluorescent reporter tags, which are more difficult to generate as chemically synthesized donors. However, cost-effective enzymatic synthesis of long ssDNA can be challenging. In this study we performed a side-by-side comparison of cssDNA produced from phagemids with lssDNA produced either using published protocols^22^ or a biotin-streptavidin capture method that we utilized^29, 64^ (Table 1). The biotin-affinity approach for making lssDNA permits the efficient synthesis of longer DNA templates and is not subject to the potential fidelity issues of RT-based approaches, as the lssDNA is generated entirely by high-fidelity DNA polymerases. Overall, we found that phagemid-derived cssDNA, when co-delivered with Cas9 or Cas12a RNPs, is highly effective in achieving targeted integration of DNA cassettes in mammalian cells. The production of cssDNA templates using phagemids is time- and cost-effective in comparison to methods for generating lssDNA donors, in part because it requires fewer electrophoretic or affinity purification steps.

We examined the relative efficacy of HDR potentiated by different ssDNA donor compositions in the context of different Cas nuclease effectors, relative strand orientations and donor doses. We initially assessed the effects of these parameters and the donor compositions on HDR efficiencies using a modified traffic light reporter system (TLR-MCV1). This fluorescence-based system permits simultaneous evaluation of imprecise and HDR-based editing efficiencies with a range of Cas9 and Cas12a effectors. While we observed robust integration of the GFP correction cassette using SpyCas9, Cas12a nucleases achieved higher overall yields of donor integration. The effects of ssDNA strand orientation, whether lssDNA or cssDNA, exhibited cell-line- and target-site-specific variability. Overall, the potency of cssDNA donors was significantly higher (i.e., effective at lower doses) than lssDNA donors, with the TLR-MCV1 reporter as well as at endogenous sites. When used in conjunction with SpyCas9 RNP, cssDNA-based HDR was robust even at concentrations as low as 1pmol cssDNA donor per 100,000 cells, while lssDNA donors were 2- to 10-fold less effective at this dose. The use of large amounts of donor DNA to drive longer insertions in cell lines typically poses toxicity issues. The improved HDR potencies of cssDNA donors relative to those of the corresponding lssDNAs could arise from higher stability of these templates in cells, since the circular topology likely confers some resistance to exonucleases. Consistent with this hypothesis, post-synthetic circularization of a lssDNA template increased the HDR efficiency by about two-fold in K562 cells to levels that were comparable to phagemid-sourced cssDNA.

In addition to exonuclease resistance conferred by circular topology, phagemid-derived ssDNA templates offer several other advantages over lssDNA templates generated using RT- or PCR-based approaches: 1) cssDNA can be generated with longer donor cassettes.^65^ Excluding the encoded bacterial and phage DNA sequences (~2,200 bp), our experience indicates that DNA cassettes up to ~10 kb (Supplementary Fig. S12) can be readily incorporated into the phagemid vector for successful ssDNA generation, without any concomitant increase in generation cost or production of truncated products. While linear ssDNA has the advantage of only containing the sequence of interest, creating donors of this length would be extremely challenging with TGIRT and potentially cumbersome even for PCR-based approaches. 2) TGIRT does not possess proofreading activity, and therefore the fidelity of ssDNA products that it produces is of concern, especially for longer donors. By contrast, the biotin-streptavidin affinity purification-based approach for generation of lssDNA and phagemid-derived cssDNA described in this paper can be used to generate accurate and full-length ssDNA. 3) The cost of generating full-length cssDNA molecules is modest compared to lssDNA generation by RT-based methods or the biotin-streptavidin affinity purification approach, which use expensive enzymes and DNA purification kits (Table 1). Moreover, the production of cssDNA can be readily scaled up to generate several micrograms of DNA at a relatively low cost, which would be cumbersome to accomplish using *in vitro* approaches. Overall, the efficacy of phagemid-derived cssDNAs as HDR templates, combined with their ease and economy of production, make them an attractive alternative for precise genome editing. cssDNA templates should prove advantageous for the efficient insertion of long DNA cassettes in a variety of different cell types and can be leveraged for basic science and potentially stem cell-based therapeutic applications.

## Methods

### Plasmids

All the plasmids generated in this study were made using standard molecular biology techniques. A list of primers used to make the donor DNA templates are listed in Supplementary Table S4. A list of plasmids created is provided in Supplementary Table S5, and plasmids have been deposited in Addgene for distribution (Deposit #75933, 75862, 87448, and 107317).

### Generation of ssDNA templates using phagemids

#### Preparation of cells

1 ml of 2xYT media with 100μg/ml ampicillin was inoculated with a colony of XL1-Blue cells transformed with the phagemid of interest. After culturing cells at 37°C for ~8 hours or until the media became slightly cloudy (OD_600_ ~0.1), 50 μl of VCSM13 phage (10^10–11^ pfu/ml) was added to the bacterial culture and incubated without shaking at RT for 20 minutes. Cells were then transferred to 250 ml 2xYT media with 100 μg/ml ampicillin and cultured at 37°C for 1-2 hours. To select for cells that had been infected by the phage, kanamycin was added to the cells to a final concentration of 75 μg/ml and cultured overnight.

#### Phage pellet preparation

Cells were pelleted from the media by centrifugation at 10,000g for 20 minutes. The supernatant containing phage was filtered through a vacuum filter (pore size 0.22 μm) to eliminate cell debris and remove any remaining bacterial cells from the supernatant. DNase I (Sigma) was added to a final concentration of 10 μg/ml and incubated at 37°C for 3 hours to eliminate any remaining dsDNA contamination in the supernatant. 10 g of PEG-8000 (Sigma) and 7.5 g of NaCl was added to 250 ml of supernatant and incubated at 4°C on ice for 1 to 2 hours to precipitate the phage. The supernatant was spun at 12,000g for 30 minutes at 4°C and the supernatant was carefully poured out and the phage pellet was retained. Care was taken to remove as much PEG solution from the bottle as possible by wiping the inner surface using Kimwipes.

#### DNA extraction

The ssDNA was extracted from the phage pellet using a modification of Purelink Midiprep columns from Life Technologies. The phage pellet was resuspended in 6 mls of 1x TE buffer. 6 ml of 4% SDS was added to the phage suspension and incubated at 70°C for 30 minutes. 6 ml of Buffer N3 or 3 M Potassium acetate (pH 5.5) was then added to the solution and spun at 12,000g for 10 minutes at room temperature. During this time, the Purelink midiprep column was equilibrated by adding 10 ml of equilibration buffer. Following column equilibration, supernatant containing cssDNA was applied to the column. The column was washed twice with 10 ml of wash solution and eluted using 5 ml of elution buffer. 3.5 ml of isopropanol or 12.5 ml of 100% ethanol was added to precipitate the DNA and incubated at −80°C for 2 hours. The solution was spun at 12,000g for 30 minutes to pellet the DNA. The DNA pellet was then washed with 5 ml 70% ethanol and allowed to air-dry. The ssDNA was then resuspended in 50-100μl of TE buffer and stored at −20°C. We typically obtain 100-200 μg of cssDNA from a 250 ml culture.

### Generation of ssDNA templates using TGIRT

Single-stranded DNA donors were generated using reverse transcription of an RNA intermediate using TGIRT-III, as previously described.^22^ Briefly, the donor sequence and its homology arms were cloned into a plasmid. Eight 50 μl PCR reactions were set up for each donor to amplify the cloned donor using forward primers that contain a 5’ overhang encoding the T7 promoter. The generated PCR products were pooled and purified using carboxylate-modified magnetic bead solution (GE Healthcare #65152105050250). The purified DNA was used to generate the corresponding RNA by *in vitro* transcription using HiScribe T7 polymerase (NEB #E2040S). After purifying the RNA with carboxylate-modified magnetic beads, the reverse transcription reaction was generated using 400 pmol of RNA, 800 pmol of reverse-transcription primer and 15 μl of 25 mM dNTP mix. After annealing the primer at 65°C for 5 minutes, then on ice for 5 minutes, 3 µl of TGIRT-III enzyme (InGex) was added and the reaction incubated at 58°C for 3 hours. The remaining RNA was hydrolyzed by base [0.5 M NaOH, 0.25 M EDTA (pH 8.0)] incubation at 95°C for 10 minutes. The NaOH was neutralized with an equal volume of 0.5 M HCl. The generated ssDNA donor was purified by carboxylate-modified magnetic beads and eluted with 20 μl or 15 µl of RNase-free water containing 2 mM Tris-HCl (pH 8.0).

### Generation of ssDNA templates using biotin and streptavidin-based affinity purification

The PCR product template for producing ssDNA was generated using one unmodified and one 5′-biotinylated primer (purchased from IDT). The High-Fidelity PCR product was purified by PCR clean-up gel extraction (QIAquick Gel Extraction Kit). Streptavidin magnetic Dynabeads (NanoLink™, catalogue number M-1002; TriLink Biotechnologies, San Diego, CA, USA) were washed and resuspended in binding solution (KilobaseBINDER™, catalogue number 60101; Invitrogen, Life Technologies) as per the manufacturer’s instructions and prepared for nucleic acid binding (17 μg of biotinylated dsDNA/mg Dynabeads, 0.8-3.3 kb). The prepared streptavidin-coated beads were incubated with biotinylated PCR product for 3 hours at room temperature or 4°C overnight while gently rotating the tubes to keep the beads in suspension. The supernatant was collected in an Eppendorf tube and biotinylated DNA-coated beads were separated with a magnet for 4 minutes. The beads were washed twice with buffer that consists of 50 mM Tris-HCl (pH 8.0), 2 M NaCl and 0.05% Tween 20 by pipetting and using a volume equivalent to the solution used for nucleic acid binding, and then the tube was placed on the magnet for 2 min to collect the beads. The beads were then washed once with buffer containing 10 mM Tris-HCl (pH 8.0) and 50 mM NaCl. The bead-containing solution was then transferred to a fresh tube and the beads were separated from the solution using a magnet for 3 minutes.

### Denaturation of dsDNA

Streptavidin beads bound to the biotinylated DNA were incubated with 155 μl of 0.1 N sodium hydroxide solution (NaOH) for 1 minute at room temperature to achieve alkaline denaturation of the biotinylated and non-biotinylated strands of the PCR product. Biotinylated ssDNA-coated beads were then separated with a magnet for 1 minute. The supernatant was then transferred to a new 1.5 ml tube and the tube was placed back on the magnetic stand for an additional 1 minute. The solution containing the non-biotinylated strand was immediately neutralized by the addition of 1 M glacial acetic acid (15 μl of 1 M glacial acetic acid to neutralize 150 μl of 0.1 N NaOH), and an equal volume of 10 mM Tris-HCl (pH 7.5) solution was then added. The sample was applied on a Spin-X centrifuge tube filter (0.22 μm cellulose acetate) to remove any beads (~0.85 μm) and transferred to a fresh tube. The non-biotinylated strand was precipitated using ethanol precipitation and then re-dissolved in water.

### Circularization of linear ssDNA

To circularize linear ssDNA donors generated by PCR using one 5′-phosphorylated and one 5′-biotinylated primer (IDT), the non-biotinylated and phosphorylated ssDNA was generated by the affinity purification method described above. Subsequently, phosphorylated ssDNA (e.g., ~20 pmol) was annealed with a 1.2-fold molar excess of splint oligonucleotide (24 pmol) that spans the two ends of the ssDNA in 1x *E. coli* DNA ligase buffer solution (NEB) to a final volume of 200 μl by heating the solution to 95°C for 2 minutes and then cooling the reaction on ice for 2 minutes. After annealing, 40 units of *E. coli* DNA ligase (NEB) was added to the solution and incubated at 45°C for 1 hour to allow ligation of the ssDNA ends to proceed to completion. The solution was then treated with 40 units of exonuclease I (NEB) and 40 units of exonuclease III (NEB) and incubated at 37°C for 30 min to eliminate linear ssDNA. Exonucleases were inactivated at 70° for 20 minutes. The cssDNA was cleaned by a NucleoSpin^®^ (Macherey-Nagel GmbH & Co. KG, Düren, Germany) column, concentrated using ethanol precipitation, and then re-dissolved in water. DNA fractions were then run on a denaturing agarose gel (2%, 70V, 2hr) to examine the integrity and purity of the cssDNA.

### Cell culture

HEK293T cells were maintained in DMEM media supplemented with 10% FBS and 1% penicillin and streptomycin (Gibco). K562 cells were maintained in RPMI 1650 media with 1 mM glutamine supplemented with 10% FBS and penicillin and streptomycin. All the cells were maintained in a humidified incubator at 37°C and 5% CO_2_.

### Electroporation of Cas9 or Cas12a RNPs

All electroporations were done using the Neon transfection system (Invitrogen). 20 pmol of SpyCas9-3xNLS, AspCas12a, LbaCas12a, or FnoCas12a protein, along with 25 pmol of sgRNA (for SpyCas9) or 60 pmol of crRNA (for Cas12a), was added per reaction. Guide RNA was either generated using *in vitro* transcription (TLR-MCV1 locus) or was purchased from Synthego (for SpyCas9 sgRNAs targeting endogenous loci). RNP and guide RNA was precomplexed in buffer R for 10-20 minutes at room temperature and the solution was made up to a final volume of 12 μl. For electroporating K562 cells, 150,000-200,000 cells per reaction were used. Cells for a reaction were spun down and the media was carefully removed. Cells were resuspended in 10 μl of buffer R containing the desired nuclease and nucleofected with 3 pulses of 1600V for 10 ms using a 10 μl Neon Tip. Cells were then plated in 24-well plates into 500 μl of RPMI 1650 media supplemented with 10% FBS and cultured in a humidified incubator at 37°C and 5% CO_2_ for 3-4 days for TLR experiments, and for 2 weeks for experiments with donors to knock in fluorescence tags at endogenous sites, before analysis of samples using flow cytometry. For all HDR experiments except those in Figure 1E, 1 pmol of cssDNA, linear ssDNA or plasmid donor DNA was used. Donor DNA was added to the cells resuspended in buffer R or buffer R containing Cas9/Cas12a RNP.

For experiments with HEK293T cells, roughly 100,000 cells per reaction were used and the cells were given 2 pulses of 1100 V for 20 ms. For experiments shown in Figure 1C and 1D, 3 pmols of cssDNA, lssDNA or plasmid donor DNA were used. For the rest of the experiments except those in Figure 1E, 1 pmol of donor DNA was used for HDR experiments.

### FACS analysis

Cells were first washed twice with 1x PBS before analysis using flow cytometry. All flow cytometry was performed on MACSQuant VYB by Miltenyi. For detection of mCherry signal, a yellow laser (wavelength 561nm) was used for excitation and a 615/20 nm emission filter was used. To detect GFP signal, a blue laser (excitation wavelength 488 nm and emission filter 525/50 nm) was used. 20,000 events were recorded for each sample and data was analyzed using Flowjo V.9.0 software. Cells were first gated on FSC-A and SSC-A plot to remove cell debris. This population was further plotted on an FSC-A vs FSC-H plot to circumscribe the single cell population. Finally, a bivariate plot between FITC-A and txRED signal was used to estimate the percentage of GFP-positive or mCherry-positive population and was reported in this study as a measure of gene editing or homologous recombination as applicable.

### TIDE analysis

Genomic DNA was extracted from mammalian cells using Sigma Genelute kit or Qiagen DNeasy Blood & Tissue Kits. PCR reactions were performed using genomic DNA as template with primers listed in Supplementary Table S4 as per the manufacturer’s directions. Subsequently, PCR product was purified using the Zymo DNA purification kit and sent for analysis by Sanger sequencing along with primers listed in Supplementary Table S4. The chromatograms were analyzed with the TIDE analysis webtool^55^ (https://tide.nki.nl/).

### Cas9 and Cas12a purification

Protein purification for 3xNLS-SpyCas9 and Cas12a-2xNLS proteins followed a common protocol as previously described.^66^ The generation and characterization of the 3xNLS-SpyCas9 and LbaCas12a-2xNLS constructs have been recently described.^52, 67, 68^ The pET21a plasmid backbone (Novagen) was used to drive the expression of a hexa-His-tagged version of each protein. The plasmid expressing 3xNLS-SpyCas9 (or each Cas12a-2xNLS) was transformed into *E. coli* Rosetta (DE3) pLysS cells (EMD Millipore) for protein production. Cells were grown at 37°C to an OD600 of ~0.2, then shifted to 18°C and induced at an OD600 of ~0.4 for 16 hours with IPTG (1 mM final concentration). Following induction, cells were pelleted by centrifugation and then resuspended with Ni^2+^-NTA buffer [20 mM Tris-HCl (pH 7.5) + 1 M NaCl + 20 mM imidazole + 1 mM TCEP] supplemented with HALT Protease Inhibitor Cocktail, EDTA-Free (100x) [ThermoFisher] and lysed with a M-110s Microfluidizer (Microfluidics) following the manufacturer’s instructions. The protein was purified from the cell lysate using Ni^2+^-NTA resin, washed with five volumes of Ni^2+^-NTA buffer and then eluted with elution buffer [20 mM Tris-HCl (pH 7.5), 500 mM NaCl, 500 mM imidazole, 10% glycerol]. The 3xNLS-SpyCas9 (or each Cas12a) protein was dialyzed overnight at 4°C in 20 mM HEPES-NaOH (pH 7.5), 500 mM NaCl, 1 mM EDTA, and 10% glycerol. Subsequently, the protein was step-dialyzed from 500 mM NaCl to 200 mM NaCl [final dialysis buffer: 20 mM HEPES-NaOH (pH 7.5), 200 mM NaCl, 1 mM EDTA, 10% glycerol]. Next, the protein was purified by cation exchange chromatography [column = 5 ml HiTrap-S; Buffer A = 20 mM HEPES-NaOH (pH 7.5) + 1 mM TCEP; Buffer B = 20 mM HEPES-NaOH (pH 7.5) + 1 M NaCl + 1 mM TCEP; flow rate = 5 ml/min; CV = column volume = 5 ml] followed by size-exclusion chromatography (SEC) on a Superdex-200 (16/60) column [isocratic size-exclusion running buffer = 20 mM HEPES-NaOH (pH 7.5), 150 mM NaCl, 1 mM TCEP for 3xNLS-SpyCas9; or 20 mM HEPES-NaOH (pH 7.5), 300 mM NaCl, 1 mM TCEP for each Cas12a-2xNLS]. The primary protein peak from the SEC was concentrated in an Ultra-15 Centrifugal Filters Ultracel-30K (Amicon) to a concentration around 100μM based on absorbance at 280nm. The purified protein quality was assessed by SDS-PAGE/Coomassie staining to be >95% pure and protein concentration was quantified with Pierce^TM^ BCA Protein Assay Kit (ThermoFisher Scientific). Protein was stored at - 80°C until further use.

### *In vitro* transcription

The DNA cassette containing the U6 promoter and the sgRNA framework for SpyCas9 was cloned from pLKO1-puro vector into pBluescript SK II+ backbone.^67^ Plasmids expressing each guide RNA from the U6 promoter were constructed by annealing oligonucleotides encoding guide RNA and cloning it into BfuAI cleavage sites in this vector (Supplementary Table S2). Templates for *in vitro* transcription (IVT) of SpyCas9 guides were amplified from the cognate plasmids using NEB Q5 High-Fidelity DNA Polymerase for 30 cycles (98°C, 15s; 65°C, 25s; 72°C, 20s) using primer sets designed to include the T7 scaffold (Supplementary Table S4). For crRNA generation for Cas12a orthologs, templates for *in vitro* transcription were generated by PCR amplification of oligonucleotides designed to include the T7 scaffold along with the guide RNA and a 15-mer overlap sequence to allow annealing between the oligos (Supplementary Table S4). The oligonucleotides encoded the full-length direct repeat crRNA sequence.^67^ Thirty cycles of amplification were conducted using NEB Q5 High-Fidelity DNA polymerase (98°C, 15s; 60°C, 25s; 72°C, 20s). The PCR products were purified using the Zymo DNA Clean & Concentrator Kit (Zymo Cat. #D4005). IVT reactions were performed using the HiScribe T7 High Yield RNA Synthesis Kit using 300 ng of PCR product as template (NEB Cat. #E2040S). After an incubation for 16 hours at 37°C, samples were treated with DNase I for 40 mins at 37°C to remove any DNA contamination. Each guide RNA was purified using the Zymo RNA Clean and Concentrator Kit. Final RNA concentration was measured using a Nanodrop instrument and stored at −80°C until further use.

### Statistical Analysis

R, a system for statistical computation and graphics, was used for the analysis.^69^ Percentage of knock-in was first arcsin-transformed to homogenize the variance. Levene’s test indicates that the assumption of homogeneity of variances was met. For Figure 2B and Supplementary Fig. S8, three-way analysis of variance (ANOVA) with Completely Randomized Design was performed to test whether there were main effects of DNA topology, target gene and fluorescent tag and whether there was a gene- or/and fluorescent tag-dependent topology effect. When no significant gene- or fluorescent tag-dependent topology effect was found, the main effect of DNA topology was reported. Otherwise, two levels of topology were compared within each combination of genetic loci and fluorescent tag under the ANOVA framework using the lsmeans package^70^ if there was a significant difference among different treatments (F-test p < 0.01). For Figure 1D, the three primary factors considered were DNA topology, Cas type and orientation. For the other figures, two-way analysis of variance (ANOVA) with Completely Randomized Design was performed to test whether there was an overall difference among different treatment groups. When the F-test was significant (p < 0.01), predefined contrasts were performed within the ANOVA framework using the lsmeans package. P values were adjusted using the Hochberg method to correct for multiple inferences.^71^

## Author Contributions

E.J.S., P.D.Z. and S.A.W. directed the study. S.I., A.M., E.J.S. and S.A.W. conceived the study. S.I., A.M., J.V.B., B.P.R., R.I., J.L., P.L., E.M. and J.S.B. performed experiments. L.J.Z. performed statistical analysis of data. S.I., A.M., E.J.S. and S.A.W. analyzed data. S.I., A.M., E.J.S. and S.A.W. wrote the manuscript with contributions from all authors.

## Acknowledgements

All new reagents described in this work are being deposited with the nonprofit plasmid-distribution service Addgene. This work was supported by US National Institutes of Health grants (1R01GM115911, Somatic Cell Genome Editing grant UG3TR002668, and 4D Nucleome grant U54DK107980) to E.J.S. and S.A.W.

## Legends for supplementary information

**Supplementary Fig. S1.**
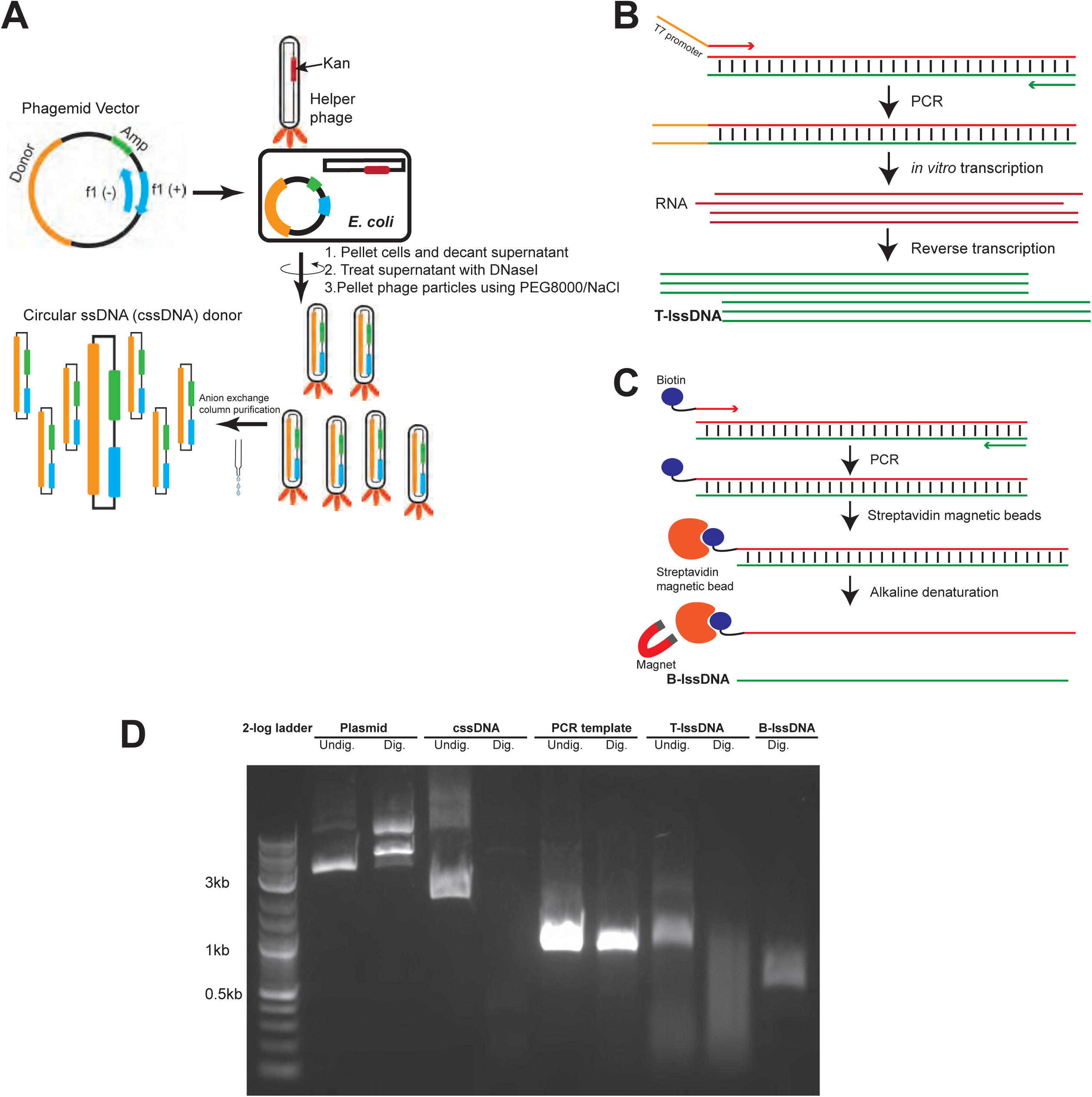
Preparation of different ssDNA templates. **(A)** Donor DNA is cloned into phagemid vectors containing an f1 bacteriophage origin of replication and an antibiotic resistance marker. The plasmid is transformed into *E. coli* cells and superinfected with a helper phage. Depending on the orientation of the f1 origin, one particular strand is packaged into phage particles and extruded into the media from which phage particles are precipitated and cssDNA is purified. (**B)** PCR product encoding donor DNA is generated using a 5’ primer containing a T7 promoter within the tail. The product is then used as a template for *in vitro* transcription to generate RNA. This RNA in turn is used as a template for reverse transcription using a reverse transcriptase such as TGIRT to generate linear ssDNA (T-lssDNA). (**C)** A PCR primer is biotinylated at the 5’ end. The resulting biotinylated PCR product is then immobilized on streptavidin magnetic beads. The immobilized PCR product is then subjected to alkaline denaturation to separate the biotinylated strand from the non-biotinylated strand. The eluted non-biotinylated DNA strand is then recovered for use as an lssDNA (B-lssDNA). **(D)** S1 nuclease digestion of DNA templates. To determine whether the templates generated are entirely single stranded, dsDNA products (Plasmid and PCR templates) and ssDNA templates (cssDNA,T-lssDNA and B-lssDNA) were digested with S1 nuclease. Undigested product (“Undig.”) was loaded alongside digested products (“Dig.”)

**Supplementary Fig. S2.**
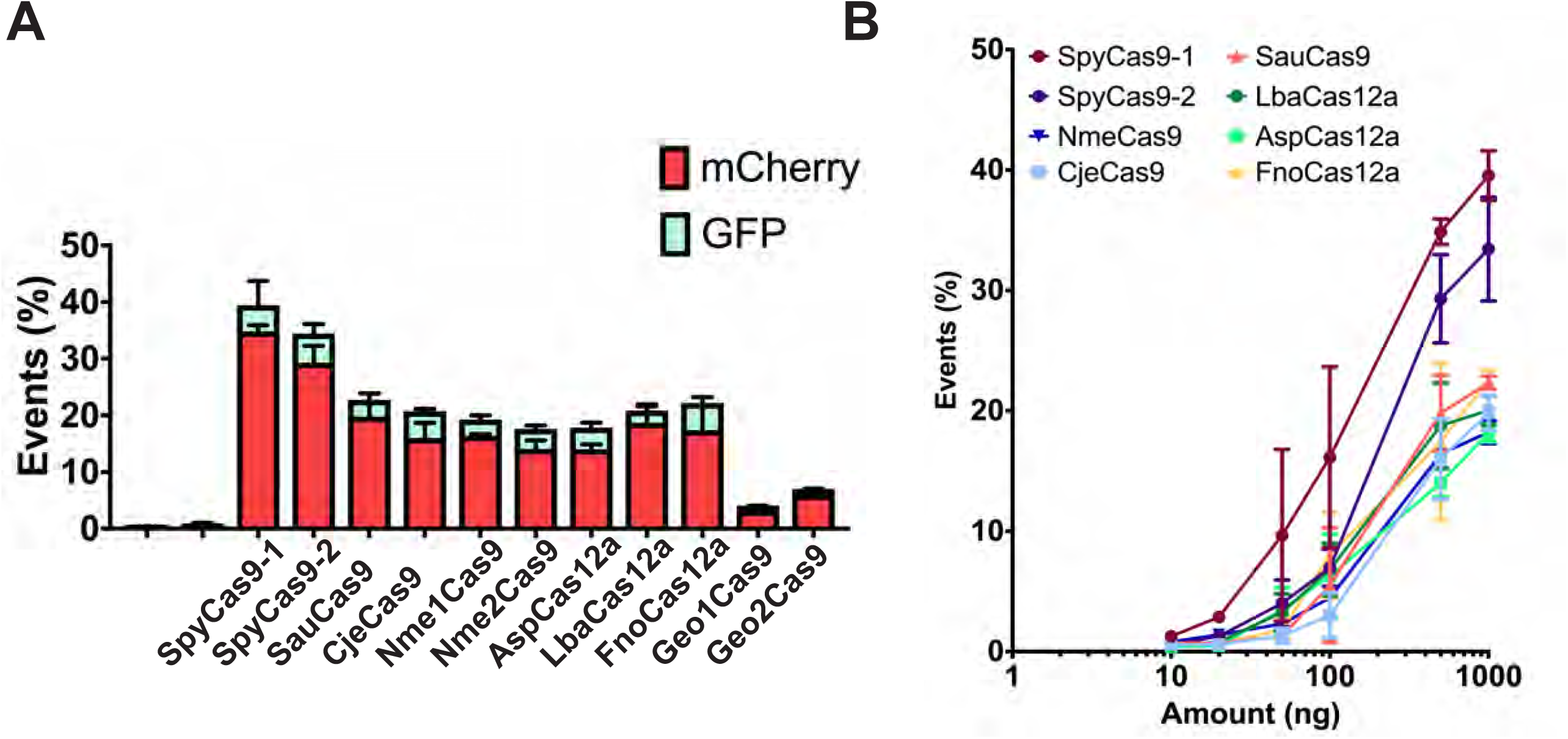
Precise and imprecise editing efficiencies for plasmid-encoded nucleases in the TLR-MCV1 reporter system. **(A)** Precise and imprecise editing efficacy of different Cas9 and Cas12a nucleases: The graph depicts the percentage of mCherry-positive (shown in red, representative of the indel efficiency) and GFP-positive (shown in cyan, representative of the HDR efficiency) cells obtained after co-delivery of 250 ng plasmid-encoded nucleases, 250 ng of gRNA plasmid and 500 ng of plasmid donor DNA template into TLR-MCV1 HEK293T cells. Bars represent the mean from three independent biological replicates and error bars represent s.e.m. **(B**) Dose dependence of editing efficiency as a function of plasmid concentration: The graph depicts the percentage of mCherry-positive cells as a function of increasing concentrations of plasmids encoding various nuclease effectors while the amount of sgRNA-expressing plasmid was held constant. Points represent the mean from three independent biological replicates and error bars represent s.e.m.

**Supplementary Fig. S3.**
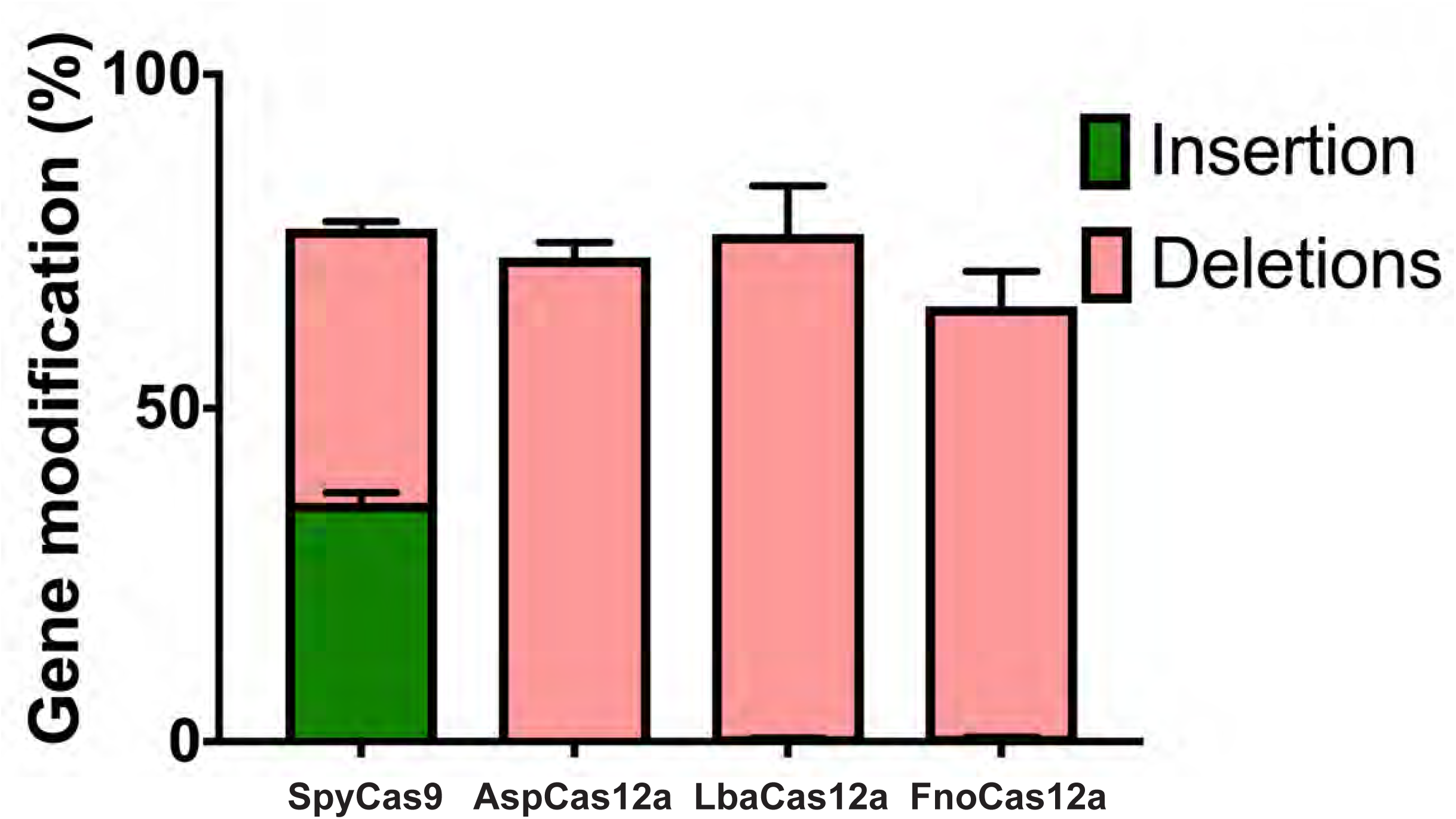
TIDE analysis to ascertain indel efficiencies at the TLR-MCV1 locus in HEK293T cells. The graph shows indel percentages observed at the TLR-MCV1 locus using SpyCas9, LbaCas12a, AspCas12a and FnoCas12a effectors based on TIDE analysis of Sanger sequencing data of the locus following nuclease treatment (in the absence of donor DNA). The green bar shows the percentage of insertions and the pink bar shows the percentage of deletions. The data show the indel percentages from three biological replicates.

**Supplementary Fig. S4.**
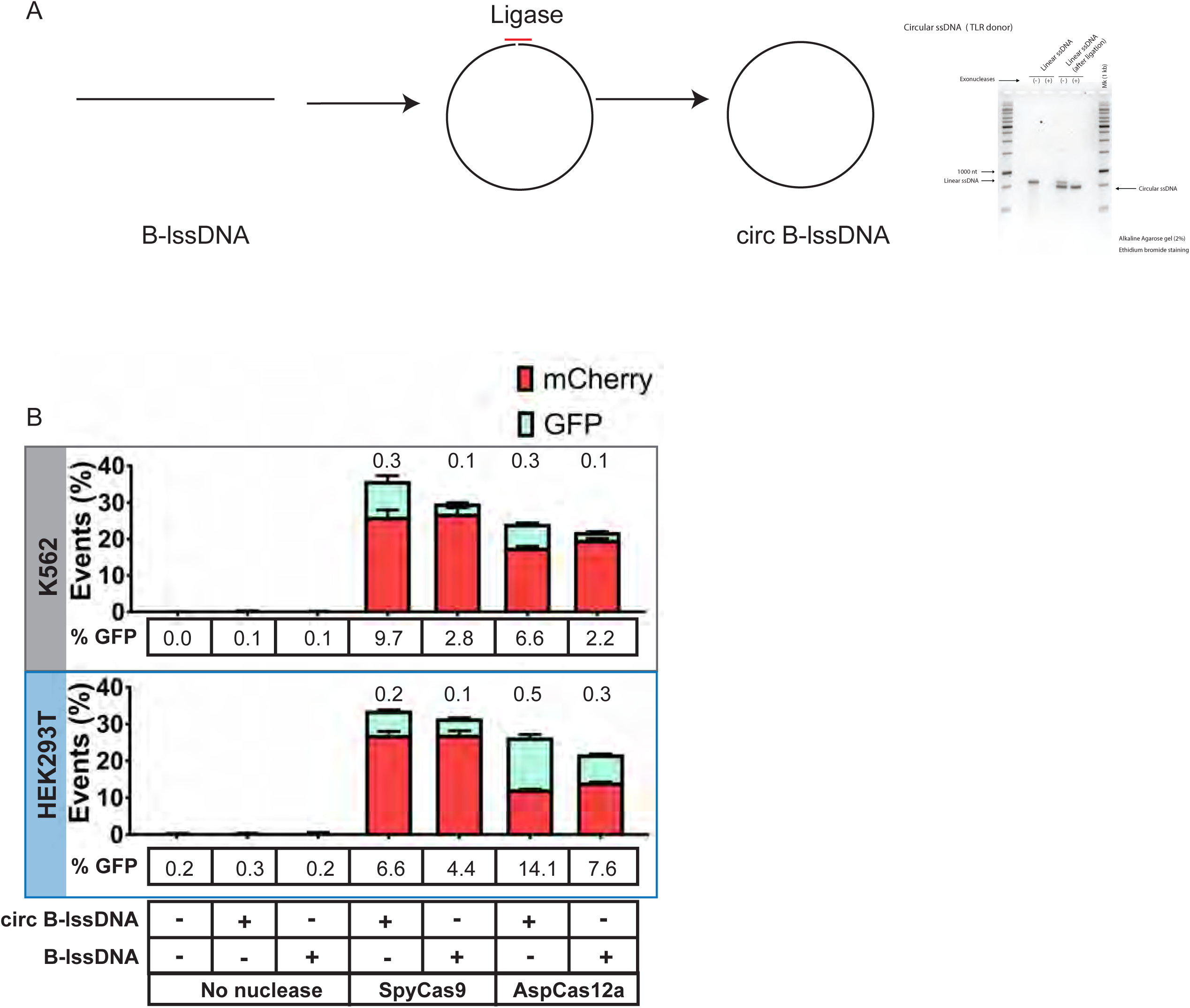
Effect of circularization of B-lssDNA on HDR efficiency. **(A)** Schematic of the approach used to generate circularized B-lssDNA. A short oligonucleotide (red) is hybridized to the B-lssDNA containing a 5’-phosphorylated end such that the oligo spans the 5’ and 3’ ends of the linear ssDNA. The sample is treated with *E. coli* DNA ligase to ligate the ends. The lssDNA sample is then treated with Exonucleases (I and III) to eliminate residual uncircularized lssDNA. The agarose gel shows unligated and ligated lssDNA before and after treatment with Exonucleases, which digest unprotected, linear DNA species. **(B)** The graph depicts the percentage of mCherry- and GFP-positive cells obtained after co-delivery of SpyCas9 with B-lssDNA and circularized B-lssDNA DNA repair templates into TLR-MCV1 K562 cells (upper grey box) and TLR-MCV1 HEK293T cells (lower blue box). Numbers above the bars indicate ratios of GFP-positive (shown in cyan) to total indels [mCherry-positive (shown in red) and GFP-positive cells]. Bars represent the mean from three independent biological replicates and error bars represent s.e.m. Numbers in the boxes below the bars show percentages of GFP-positive cells.

**Supplementary Fig. S5.**
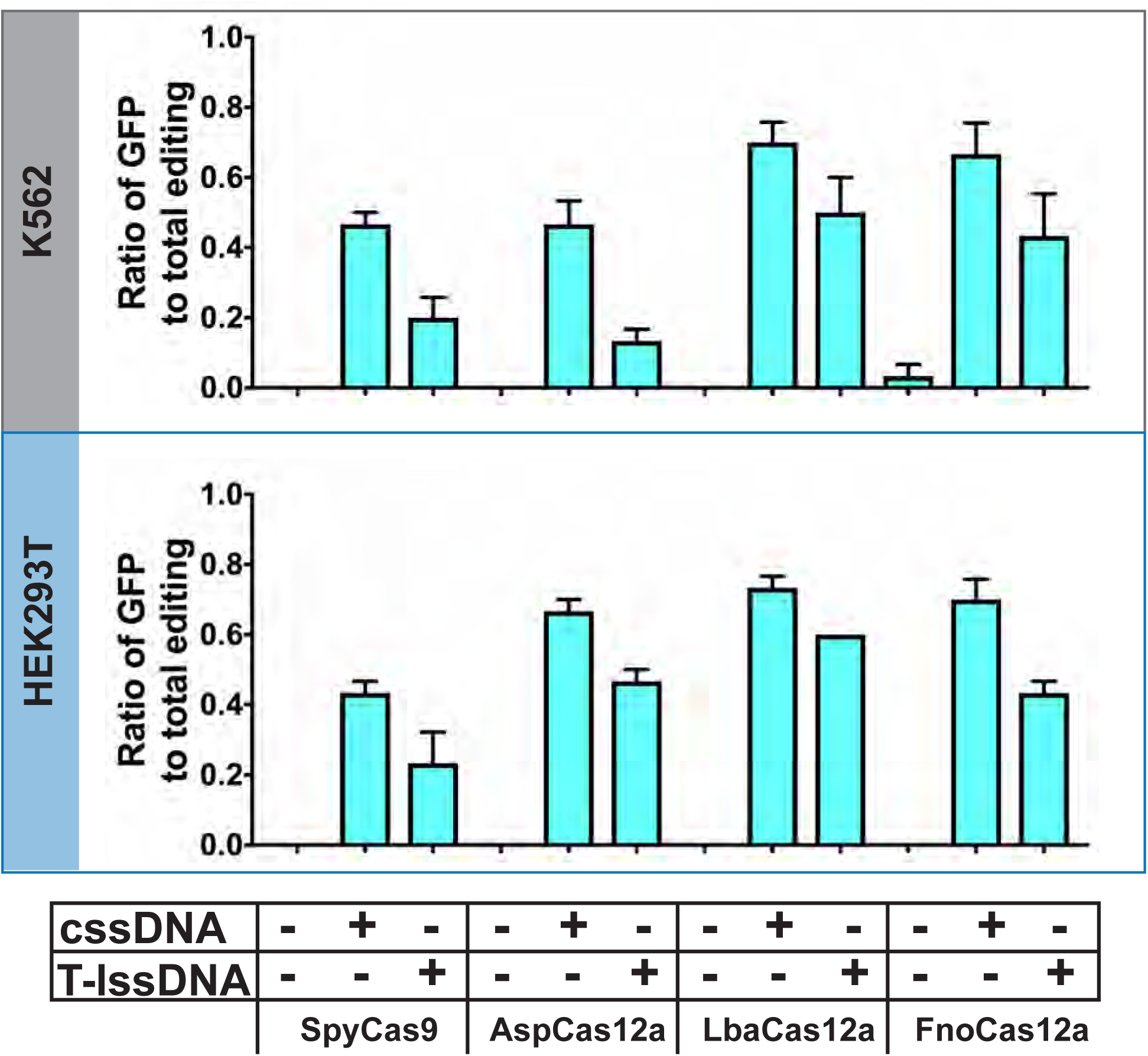
The ratio of GFP-positive cells to total editing in the samples shown in Figure 1C. The bar graph of the ratio of GFP-positive cells over total edited cells (mCherry-positive + GFP-positive cells) obtained upon treatment of TLR-MCV1 K562 cells (upper panel) and TLR-MCV1 HEK293T cells (lower panel) with SpyCas9, AspCas12a, LbaCas12a, or FnoCas12a in the absence of donor DNA or the presence of the indicated donor. Bars represent the mean from three independent biological replicates and error bars represent s.e.m.

**Supplementary Fig. S6.**
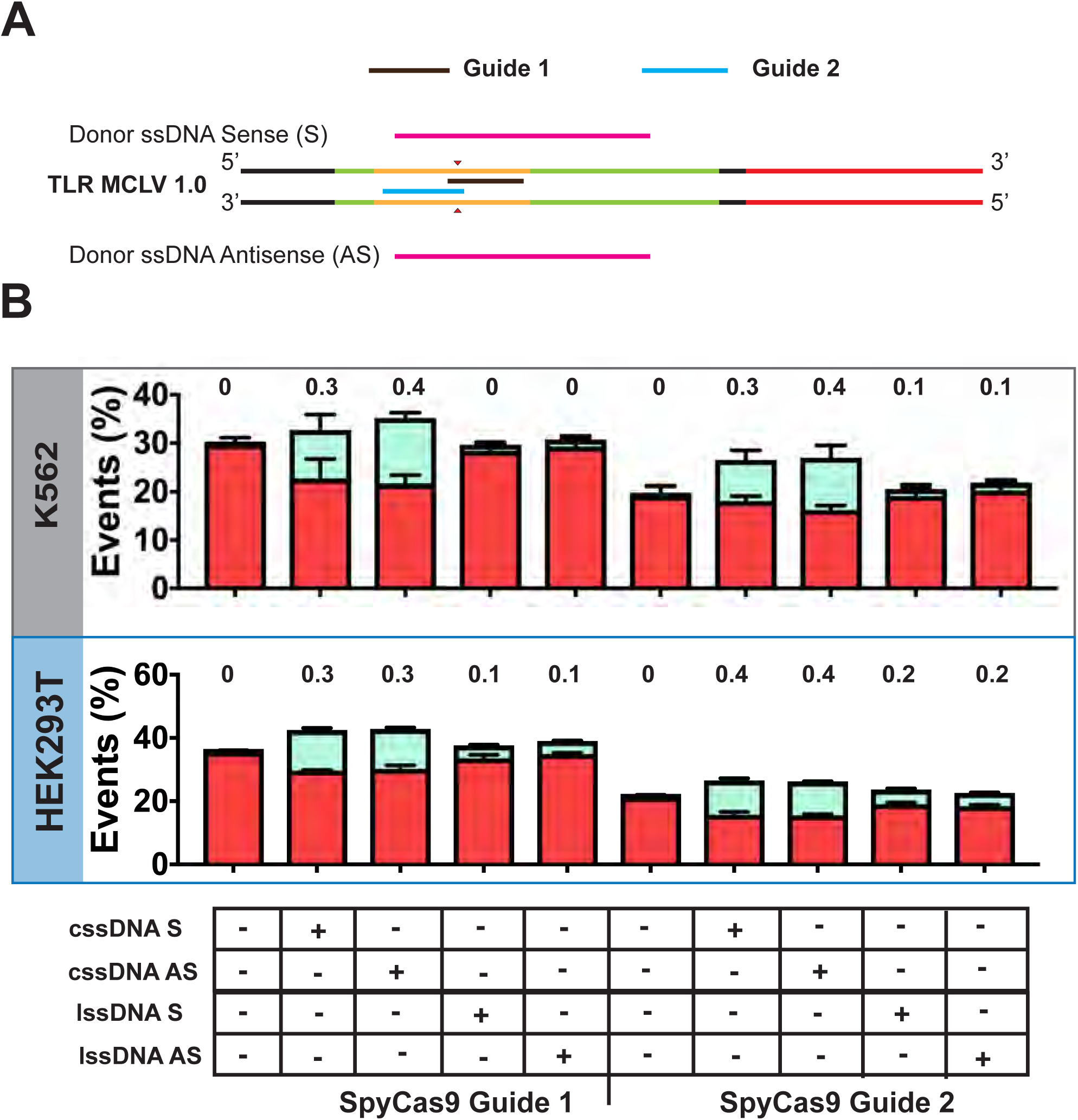
Strand dependence of guide and HDR template on knock-in efficiency. **(A)** Schematic of guide (depicted by black and blue lines) and strand orientation relative to the TLR-MCV1 target site. The magenta carrots indicate the position of the SpyCas9 DSB. The green and red lines indicate GFP and mCherry encoding regions, respectively. The orange region depicts the small insertion containing target sites for Cas9 and Cas12a proteins. **(B)** The graph depicts the percentage of mCherry- and GFP-positive cells obtained after co-delivery of SpyCas9 complexed with guides (SpyCas9 RNP) targeting either strand of the TLR-MCV1 reporter along with DNA repair templates complementary to the antisense or sense strand in K562 cells (upper grey box) and HEK293T cells (lower blue box). Numbers above the bars indicate ratios of GFP-positive (shown in cyan) to total indel events [mCherry-positive (shown in red) cells and GFP-positive cells]. Bars represent the mean from three independent biological replicates and error bars represent s.e.m.

**Supplementary Fig. S7.**
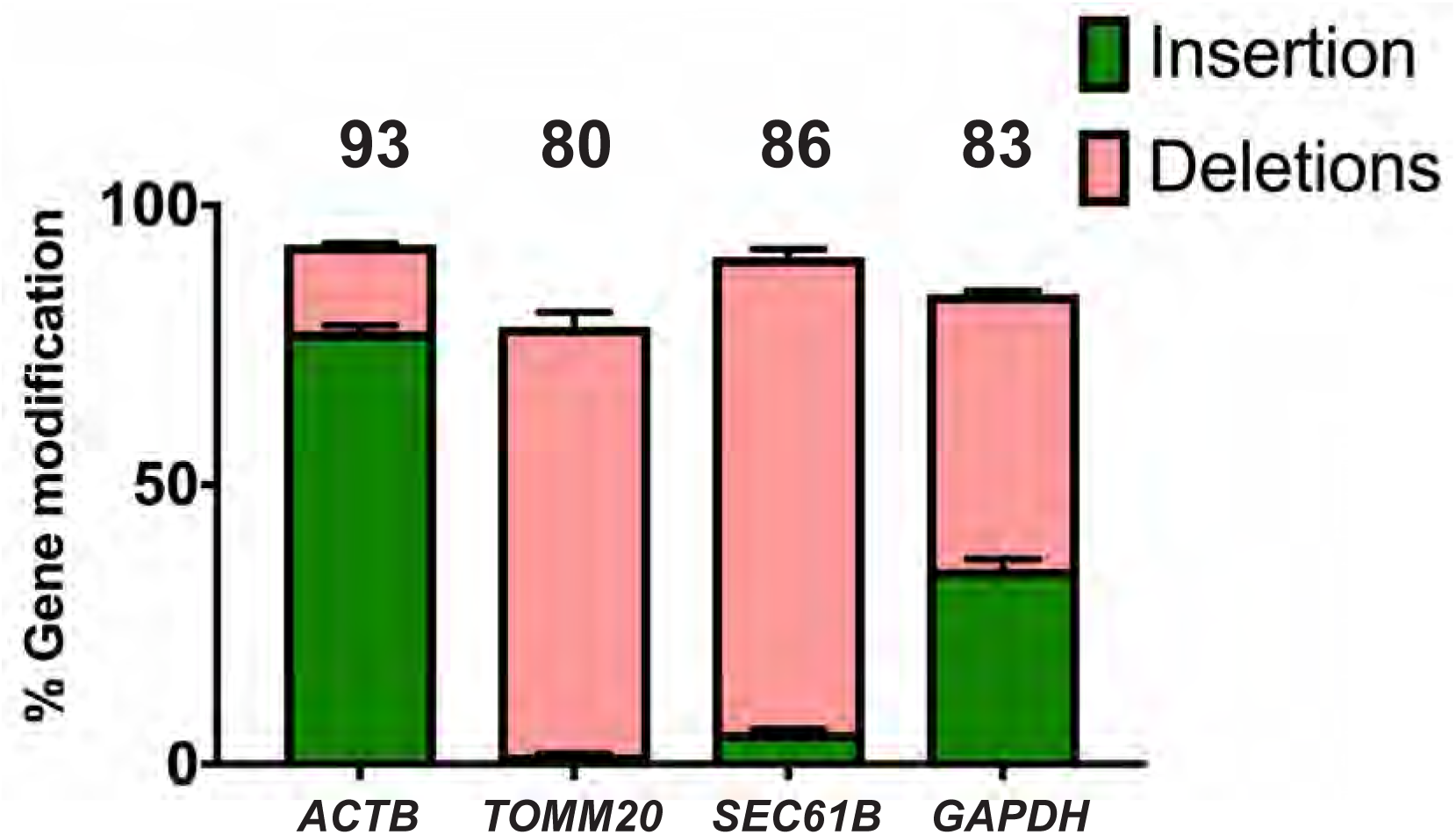
SpyCas9 gene editing efficiency at the *ACTB*, *TOMM20*, *SEC61B* and *GAPDH* loci. Genome editing was achieved by electroporation of 20 pmoles SpyCas9 complexed with 25 pmoles of guide RNA into HEK293T cells in the absence of HDR donor. The editing percentages were calculated by TIDE analysis (indicated above the bars). Pink bars indicate the proportion of deletions and green bars indicate the proportion of insertions within the indel population.

**Supplementary Fig. S8.**
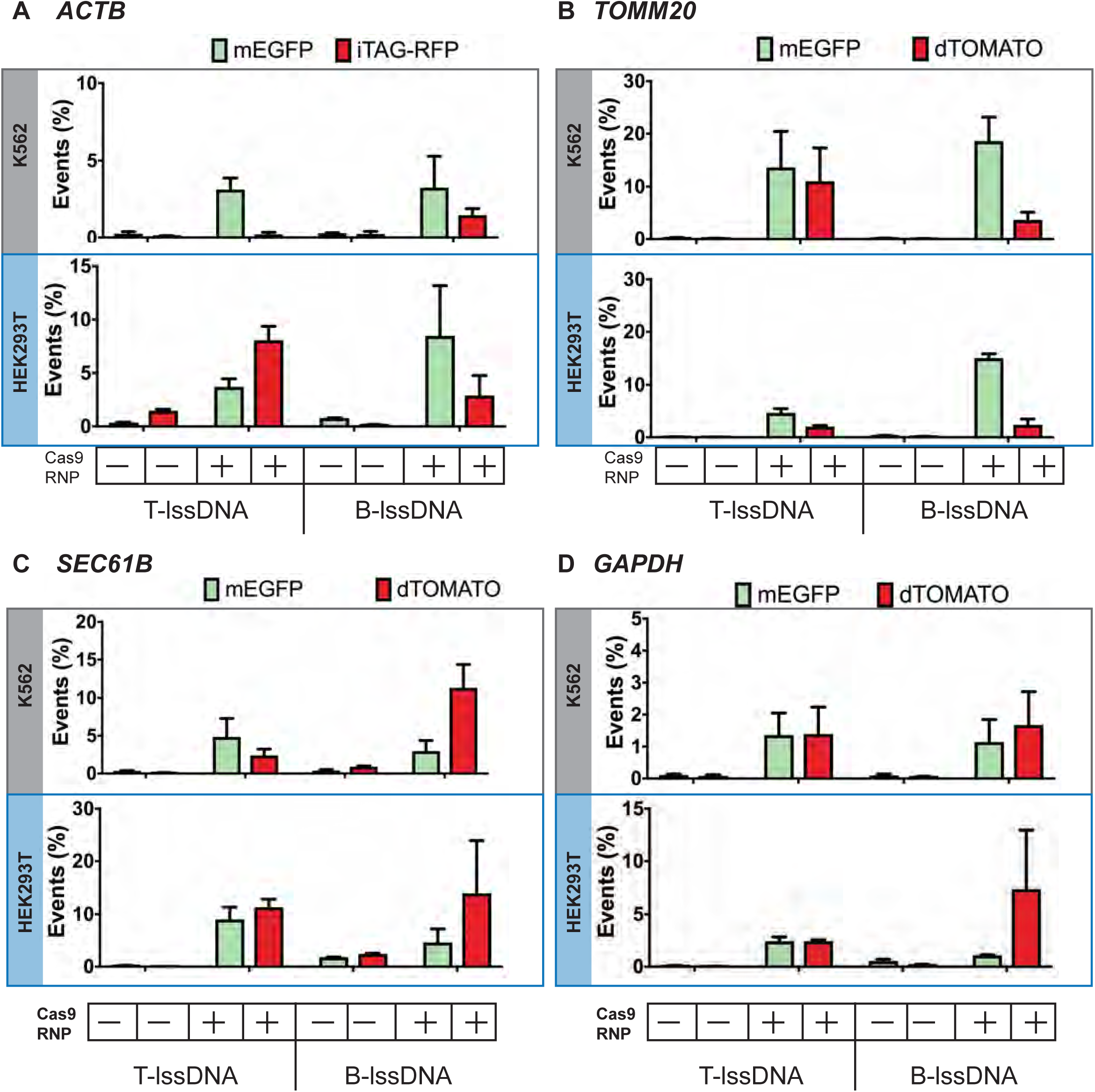
Efficiencies of fluorescent tag integration achieved with lssDNA donors generated using the TGIRT-mediated RT-PCR (T-lssDNA) or biotin-streptavidin affinity purification (B-lssDNA) approaches. Editing efficiencies for SpyCas9 RNPs and lssDNA donor delivery targeting the **(A)** *ACTB*, **(B)** *TOMM20*, **(C)** *SEC61B*, and **(D)** *GAPDH* loci in K562 (top panel) and HEK293T (bottom panel) cells are shown. Bars represent the mean from three independent biological replicates and error bars represent s.e.m.

**Supplementary Fig. S9.**
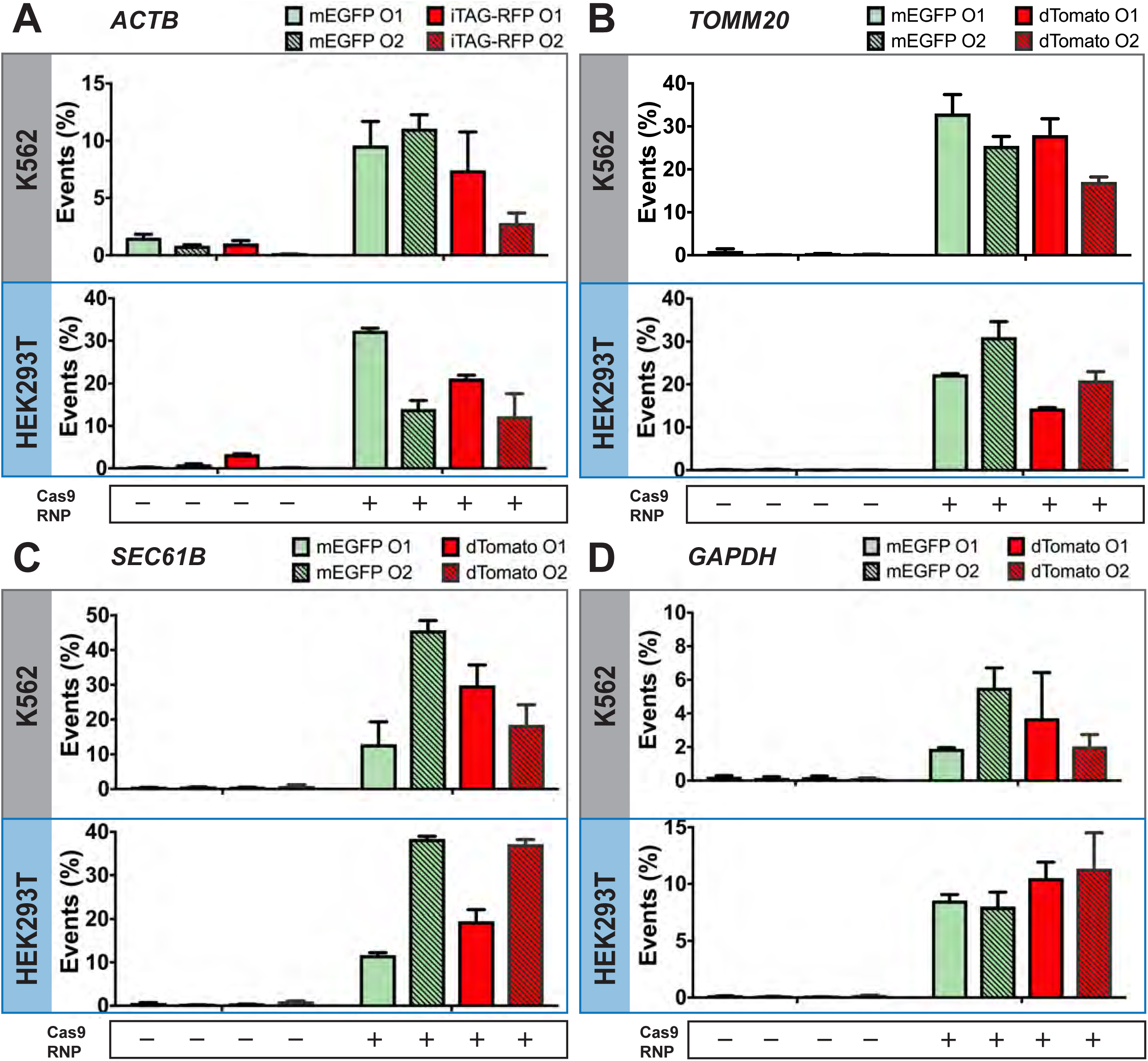
Effect of orientation of cssDNA on HDR efficiencies at endogenous loci. Editing efficiencies for SpyCas9 RNPs and cssDNA donor delivery targeting the **(A)** *ACTB*, **(B)** *TOMM20*, **(C)** *SEC61B* and **(D)** *GAPDH* loci in K562 cells (top panel) and HEK293T cells (bottom panel) are shown. Green bars indicate percentages of cells expressing GFP and red bars correspond to iTAG-RFP/dTomato integration events. Solid bars correspond to donor DNA in orientation 1 (ssDNA complementary to the antisense strand of the target gene) and hashed bars correspond to orientation 2 (ssDNA complementary to the sense strand of the target gene). Bars represent the mean from three independent biological replicates and error bars represent s.e.m.

**Supplementary Fig. S10.**
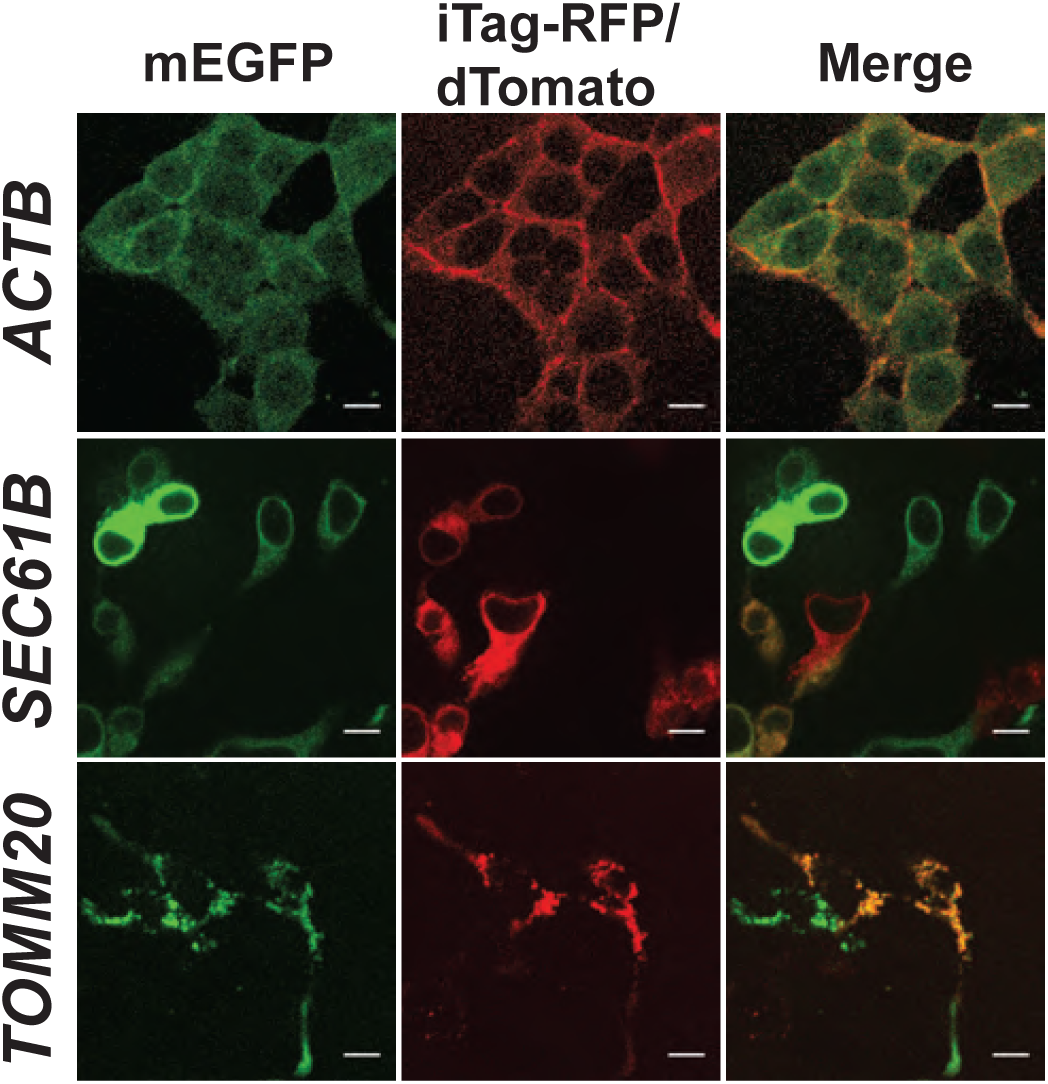
Confocal images showing tagging of GFP and iTAG-RFP at the *ACTB* locus (top panel) and GFP and dTomato at the *TOMM20* (middle panel) or *SEC61B* (bottom panel) loci in HEK293T cells from experiments shown in Figure 3.

**Supplementary Fig. S11.**
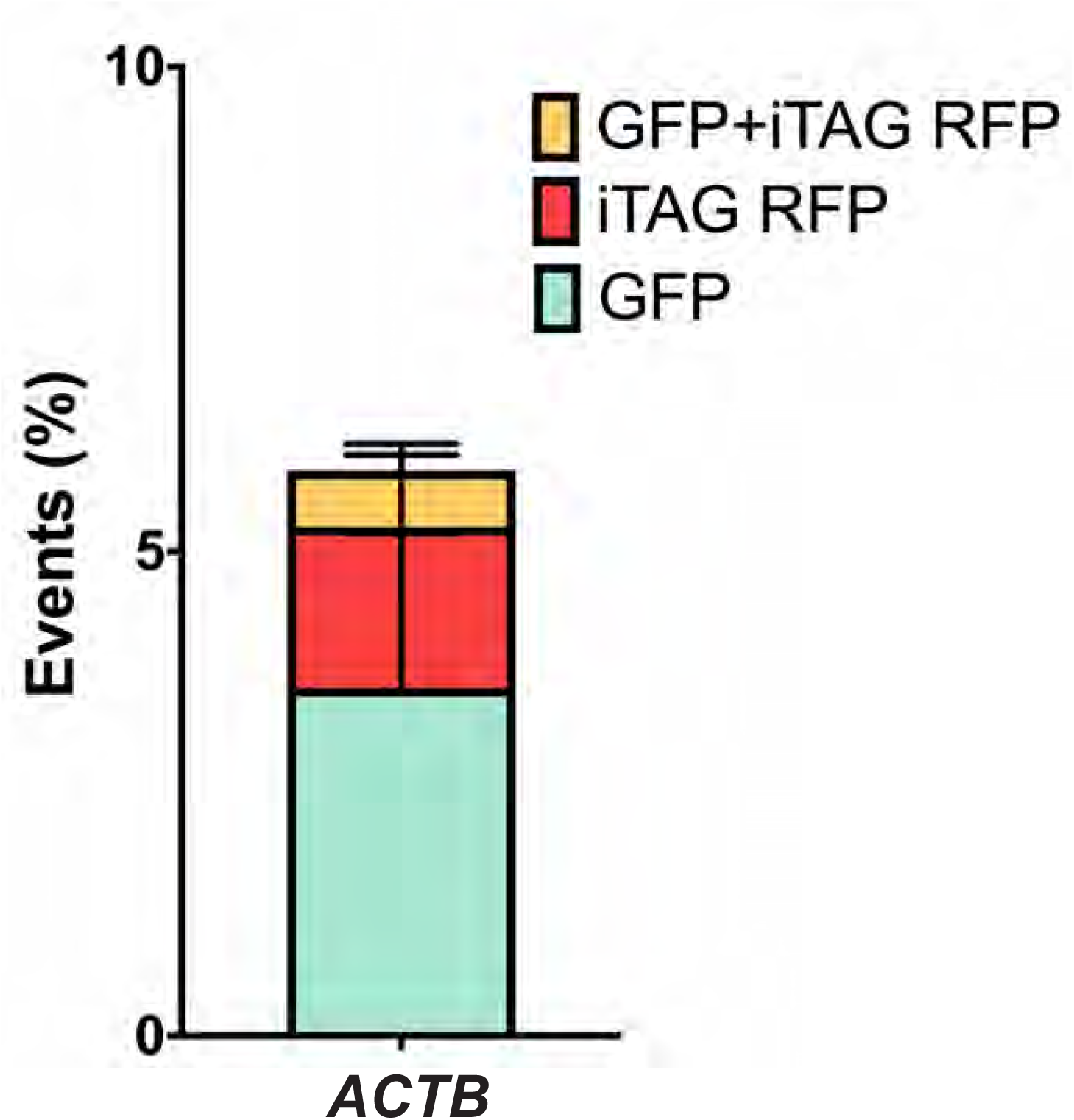
Biallelic integration of GFP and iTagRFP in K562 cells using cssDNA template. K562 cells were electroporated with 1 pmol each of GFP- and iTagRFP-encoding cssDNA templates along with 20 pmols of SpyCas9 complexed with 25 pmols of guide RNA targeting *ACTB*. Green bars represent the percentage of GFP-positive cells, red bars represent iTagRFP-expressing cells and yellow bars represents cells expressing both GFP and iTagRFP. Bars represent the mean from three independent biological replicates and error bars represent s.e.m.

**Supplementary Fig. S12.**
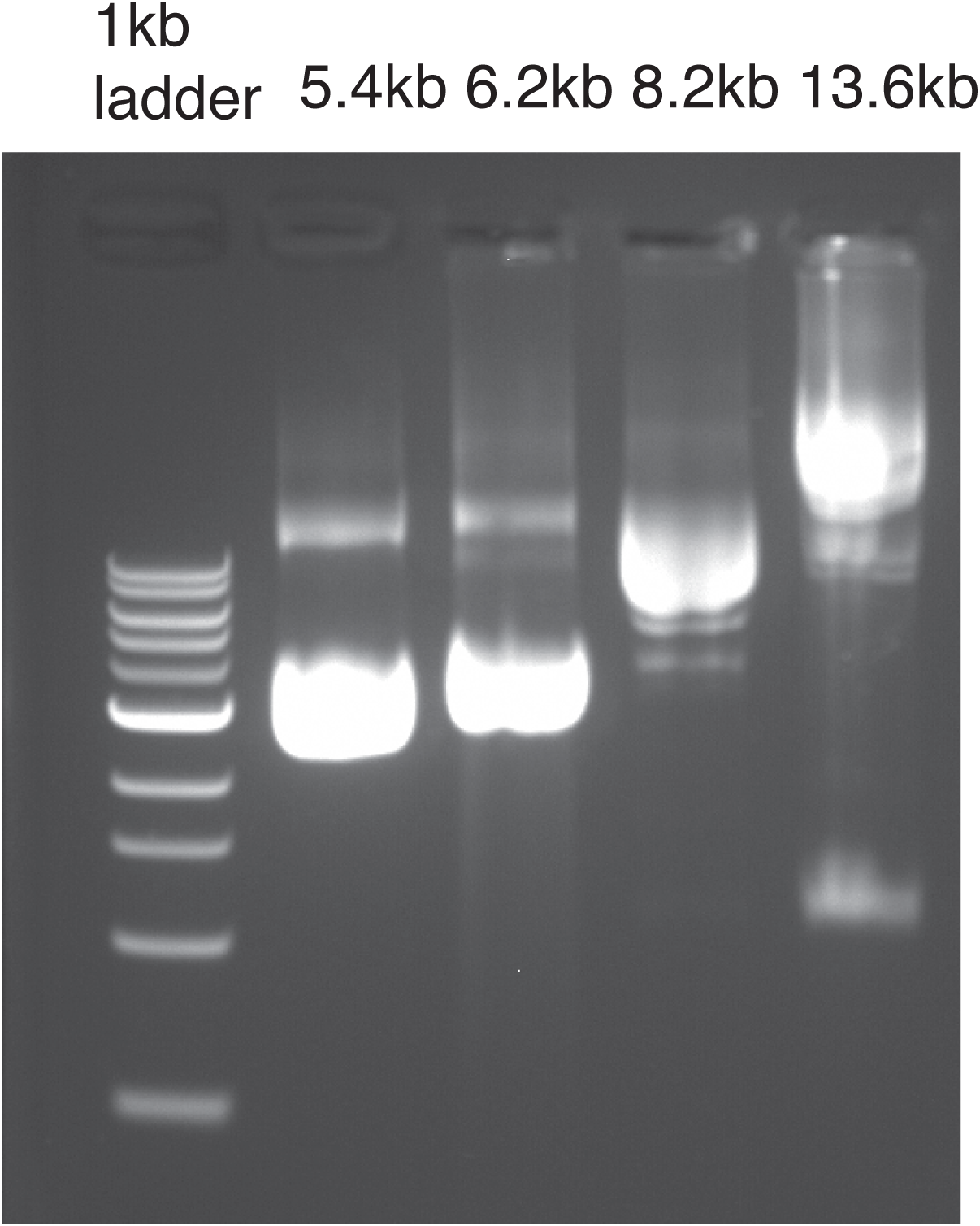
1% agarose gel image showing 1kb ladder (lane 1), as well as cssDNA generated from plasmids that are 5.4kb (lane 2), 6.2kb (lane 3), 8.2kb (lane 4) and 13.6 kb (lane 5) in length.

